# A proteostasis clock tunes bacterial dormancy by timing replication initiation

**DOI:** 10.64898/2026.03.09.710435

**Authors:** Fa-Zhan Wang, Yi-Wen Zhang, Jia-Feng Liu

## Abstract

Bacterial dormancy is a major contributor to antibiotic tolerance and persistent infection, yet the lag time before growth resumption is commonly viewed as a passive consequence of cellular recovery and remains difficult to manipulate. Protein aggregation has been linked to bacterial dormancy, but whether aggregates merely reflect cellular damage or directly regulate lag time remains unclear. Here we show that lag time is quantitatively set by cellular proteostasis through control of replication initiation. Microscopy and proteomic analyses revealed delayed aggregate disassembly as a shared proteostasis defect of long-lag bacteria. We identified the replication initiator DnaA in aggregates, whose spatial separation from the nucleoid directly suppresses DNA synthesis. Combined *in vitro* reconstitution revealed that aggregation-prone proteins drive dose-dependent, synergistic co-aggregation of DnaA, coupling aggregate burden to DnaA sequestration and lag-time extension. Across diverse genetic and physiological models, we observed a universal scaling relationship between lag-time extension and proteostasis decline. Genetic and chemical perturbations of this proteostasis-replication axis bidirectionally modulate lag time, with increasing proteostasis burden prolonging dormancy and restoring proteostasis or DnaA availability accelerating awakening. We therefore propose that proteostasis functions as a tunable timekeeping system for bacterial dormancy by regulating replication initiation, providing a basis for manipulating bacterial dormancy and antibiotic tolerance.

## INTRODUCTION

The timing of replication initiation is critical for life to balance stress tolerance and proliferation^1^. Dormant bacteria exhibit an extended lag time before resuming growth, even upon transition to nutrient-rich environments^2,3^. This delay in growth protects bacteria against antibiotics targeting actively proliferating cells, a phenomenon known as antibiotic tolerance^2–5^. *In vitro* evolution experiments reveal that bacterial lag time can be optimized to match different antibiotic treatment durations^2^, indicating that lag time is a highly evolvable trait for bacterial adaptation to fluctuating environments. Antibiotic tolerance conferred by lag extension impedes the clearance of pathogens and facilitates the subsequent evolution of resistance^3,6^. Thus, a better understanding of the molecular mechanisms that regulate bacterial lag time is essential for developing strategies to eliminate these antibiotic persisters.

Despite numerous mutations that extend lag time having been identified in bacterial toxin-antitoxin systems^7^, aminoacyl-tRNA synthetases^2,3,5^, transcriptional regulators^8^, and metabolic enzymes^9^, it has been a long-standing challenge to determine whether a common mechanism underlies these different dormant states. Recent evidence has established a correlation between bacterial exit from dormancy and disassembly of protein aggregates^4,10–12^. Nevertheless, the mechanism by which protein aggregation prevents reproduction remains poorly defined. Although various proteins associated with translation, replication, and energy generation^4,10,12,13^ have been observed in aggregates, direct evidence linking the aggregation state of a specific protein to the regulation of the corresponding pathway and bacterial lag time remains lacking.

Here, we present *in vivo* and *in vitro* evidence that sequestration of the replication initiator DnaA within protein aggregates provides a common downstream mechanism through which distinct proteotoxic stresses delay replication initiation and extend bacterial lag time. Across diverse dormant states, lag-time extension scales with the extent of proteostasis decline and DnaA sequestration. Proteostasis impairment renders dormant bacteria collaterally sensitive to proteotoxic stress, revealing a shared vulnerability of antibiotic-tolerant populations.

## RESULTS

### A diminished capacity to maintain proteostasis in dormant mutants

To understand the molecular mechanisms governing bacterial lag time, we first performed evolution experiments to identify genetic factors that shape the lag time distribution. Based on previous observations that cyclic ampicillin exposure selects for extended lag time^2^, we evolved *Escherichia coli* KLY under cyclic treatment with the antibiotic ertapenem (ETP). This antibiotic was specifically chosen for the lack of ETP-resistant carbapenemase genes in the KLY strain^2^, thus preventing the rapid emergence of resistance and enabling long-term tracking of dormancy evolution. Using ScanLag^14^ to quantify single-cell lag time, we observed that, following cyclic ETP treatments, both the mean and variance of colony appearance times increased across three parallel batch cultures (Extended Data Fig. 1a, b). Colonies exhibiting different lag times were isolated from the evolved lines and analysed by whole-genome sequencing (WGS). Mutations were identified in several aminoacyl-tRNA synthetases (*pheT*, *alaS*, *leuS*) and the cysteine tRNA (*cysT*), each conferring a specific increase in lag time (Fig. 1a, Extended Data Fig. 1c and Supplementary Table 1). In line with prior reports implicating multiple aminoacyl-tRNA synthetases in lag regulation^2,3,5,15^, these results hinted at a conserved yet unresolved link between this gene family and dormancy control.

**Fig. 1.**
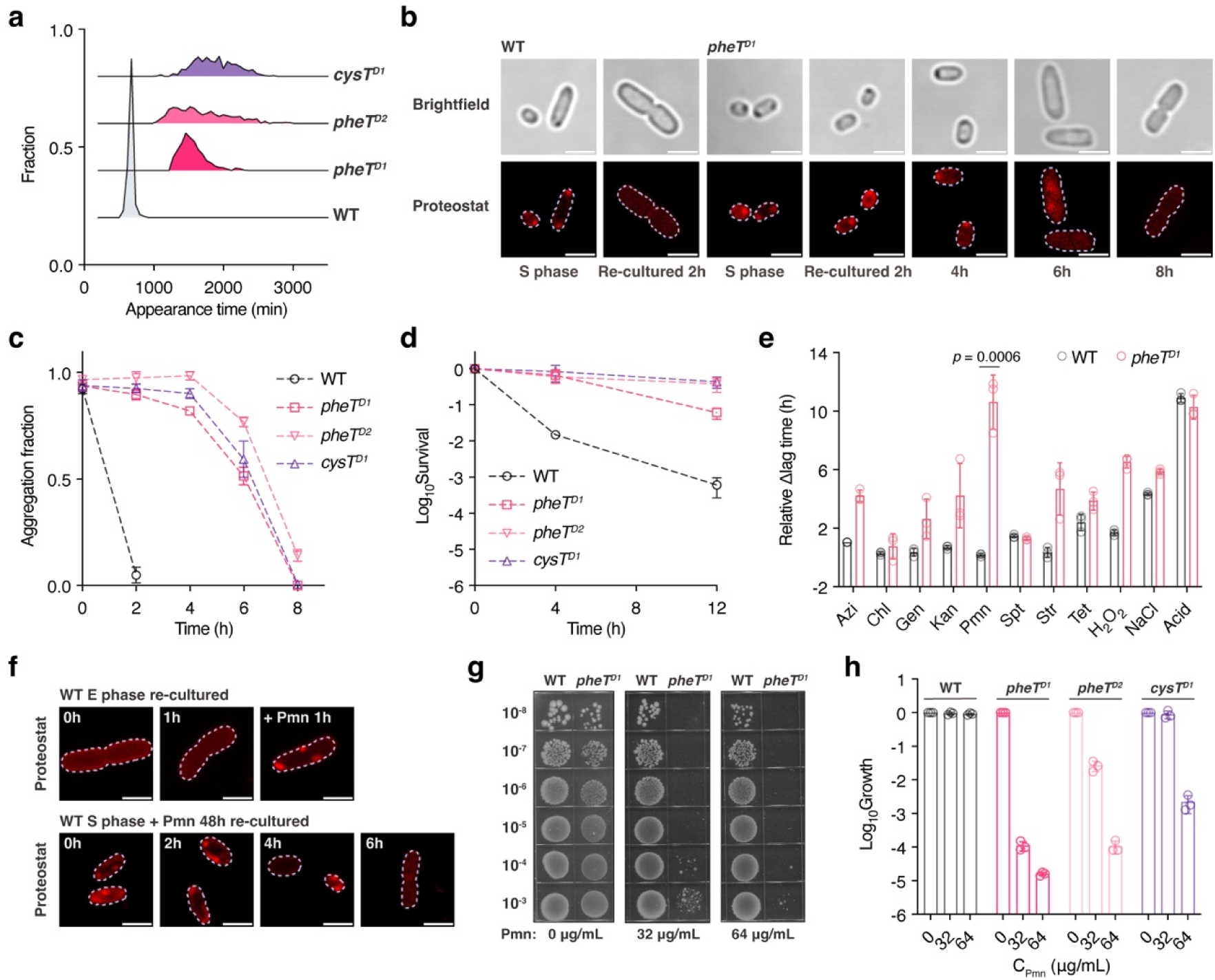
A diminished capacity of long-lag mutants to maintain proteostasis. **a,** Appearance time distribution for WT, *pheT^D1^*, *pheT^D2^*, and *cysT*^D1^ cells measured by ScanLag. Bacteria were sampled from 48-h cultures. The y-axis represents the proportion of colony-forming units (CFUs) detected at each time point. Sample sizes N = 989, 810, 678, and 457, respectively. **b**, Representative brightfield and fluorescence images of WT and *pheT^D1^* cells stained with Proteostat. Cells from the stationary phase (S phase) were re-cultured in fresh LB medium and sampled at the indicated time points. Scale bars, 2 μm. **c**, Quantification of the fraction of WT, *pheT^D1^*, *pheT^D2^*, and *cysT*^D1^ cells containing Proteostat-positive aggregates upon re-culturing in fresh LB medium. Data are presented as mean ± SD (n = 4 biological replicates). **d**, Survival of WT, *pheT^D1^*, *pheT^D2^*, and *cysT*^D1^ cells after 4h or 12h of 1 µg/mL ertapenem treatment. Data are presented as mean ± SD (n = 3 biological replicates). **e,** Effect of various stresses on the lag time of WT and *pheT^D1^*cells. Relative lag time was determined as the last time point at which ΔOD_600_ remained below 0.02, indicating minimal or no detectable growth. Relative Δlag time was normalized to the no-stress condition. Azithromycin (Azi), 4 μg/mL; Chloramphenicol (Chl), 32 μg/mL; Gentamicin (Gen), 2 μg/mL; Kanamycin (Kan), 4 μg/mL; Puromycin (Pmn), 16 μg/mL; Spectinomycin (Spt), 16 mM; Streptomycin (Str), 4 μg/mL; Tetracycline (Tet), 1 μg/mL; H_2_O_2_, 0.2 mM; NaCl, 600 mM; Acid, pH = 4. Data are presented as mean ± SD (n = 3 biological replicates). *P* values were calculated using unpaired two-tailed t-tests. **f**, Representative fluorescence images of stress-treated WT cells stained with Proteostat. Top, WT cells from the exponential phase (E phase) were re-cultured in LB or LB supplemented with 256 µg/mL puromycin (Pmn). Bottom, WT cells from the S phase were incubated in LB supplemented with 256 µg/mL Pmn for 48h and then transferred to fresh LB without antibiotics. Scale bars, 2 μm. **g**, Effect of puromycin on the growth of WT and *pheT^D1^* cells on solid plates. Serial dilutions of WT and *pheT^D1^*cells were spotted in 10-µL volumes onto plates containing the indicated concentrations of puromycin. **h**, Relative colony-forming fraction of WT, *pheT^D1^*, *pheT^D2^*, and *cysT*^D1^ cells on solid plates containing the indicated concentrations of puromycin, normalized to drug-free plates. Data are presented as mean ± SD (n = 3 biological replicates).

Notably, two independent mutations were mapped to the phenylalanyl-tRNA synthetase β subunit (*pheT*), both of which produced pronounced lag extension. Restoring the wild-type (WT) allele of *pheT* fully reverted this phenotype (Extended Data Fig. 1d). Recent studies suggest that protein aggregation could facilitate bacterial dormancy, and aggregate disassembly is critical for growth resumption^4,11,12^. To investigate the aggregation dynamics in bacteria, we stained WT and two *pheT* mutants with Proteostat, a fluorescent dye that specifically binds to insoluble protein aggregates^16,17^. Consistent with previous studies^4^, we observed the formation of insoluble protein aggregates in stationary-phase bacteria, visible as dark foci in brightfield microscopy and elevated Proteostat fluorescence (Fig. 1b and Extended Data Fig. 1e). However, quantitative analysis revealed that the proportion of cells containing protein aggregates (Fig. 1c) and the intensity of Proteostat fluorescence (Extended Data Fig. 1f) were indistinguishable between stationary-phase WT and two *pheT* mutants, despite their markedly different lag times (Fig. 1a). Subsequent isolation and quantification of insoluble protein fractions^18^ further confirmed comparable levels of aggregated proteins in these strains (Extended Data Fig. 1g). Thus, the aggregate amount in stationary-phase cells was insufficient to account for the extended lag time of the *pheT* mutants.

Intriguingly, upon nutrient replenishment, we observed that WT cells exhibited rapid protein disaggregation within two hours, whereas protein aggregates in two *pheT* mutants persisted for more than six hours until cell division (Fig. 1b, c and Extended Data Fig. 1e, f). Thus, despite similar aggregate amounts, aggregates disassembled more slowly in the *pheT* mutants before growth resumption. This temporal coupling between dormancy exit and aggregate disassembly was not restricted to the *pheT* mutants, as similar phenomena were observed in the cysteine tRNA mutant, *cysT^D1^* (Fig. 1c and Extended Data Fig. 1h). These observations identify delayed aggregate disassembly, rather than increased aggregate abundance, as a common feature of these dormant mutants.

We next determined whether the delayed aggregate disassembly is merely an epiphenomenon of extended lag time or instead reflects a compromised cellular capacity to clear protein aggregates. Given that cells with disrupted proteostasis typically exhibit reduced ability to cope with additional proteotoxic stress^19–21^, we compared the sensitivity of WT and *pheT^D1^* cells to various types of stress. Extended lag times enabled dormant mutants to survive better under ertapenem treatment (Fig. 1d). In contrast, the *pheT^D1^* mutant showed greater growth inhibition than WT cells upon exposure to puromycin (Fig. 1e and Extended Data Fig. 2a), an aminoacyl-tRNA analogue that promotes misfolded protein accumulation^22,23^. Proteostat staining revealed that puromycin treatment induced protein aggregation in fast-growing *E. coli* cells and inhibited aggregate disassembly upon dormancy exit (Fig. 1f). Compared to WT cells, puromycin treatment significantly reduced the plating efficiency of all long-lag mutants (Fig. 1g, h), indicating a compromised ability of dormant bacteria to cope with protein aggregation. Similar results were obtained upon the introduction of another well-known proteotoxic stressor, heat shock^4^ (Extended Data Fig. 2b, c).

Taken together, these findings indicate impaired proteostasis as a shared signature of dormant mutants, manifested as internal delayed aggregate disassembly and increased vulnerability to external proteotoxic stress.

### Increased PheS aggregation disrupts proteostasis and extends lag time

To identify the key components governing aggregate disassembly rate, we compared the composition of insoluble aggregates isolated from stationary-phase WT and two *pheT* mutants. Label-free quantitative proteomics identified phenylalanyl-tRNA synthetase α subunit (PheS) as a top-enriched protein within the aggregates of two *pheT* mutants relative to WT cells (Fig. 2a, b and Extended Data Fig. 3a). In *E. coli*, phenylalanyl-tRNA synthetase functions as a heterotetramer of two α (PheS) and two β (PheT) subunits, both of which are essential for bacterial viability^24^. Quantitative analysis showed that total PheS levels did not increase in the *pheT* mutants (Extended Data Fig. 3b), indicating that the increased aggregation reflects reduced PheS solubility. To validate these proteomic data, we tagged PheS with ECFP and monitored its spatiotemporal localization during dormancy. We observed the formation of PheS-ECFP aggregates in stationary-phase cells that colocalized with Proteostat signals (Fig. 2c and Extended Data Fig. 3c). Upon nutrient replenishment, these aggregates disassembled rapidly in WT cells but persisted longer in the *pheT* mutants (Fig. 2c and Extended Data Fig. 3c). These results indicate that the *pheT* mutations cause increased accumulation and delayed disassembly of PheS aggregates in dormant bacteria.

**Fig. 2.**
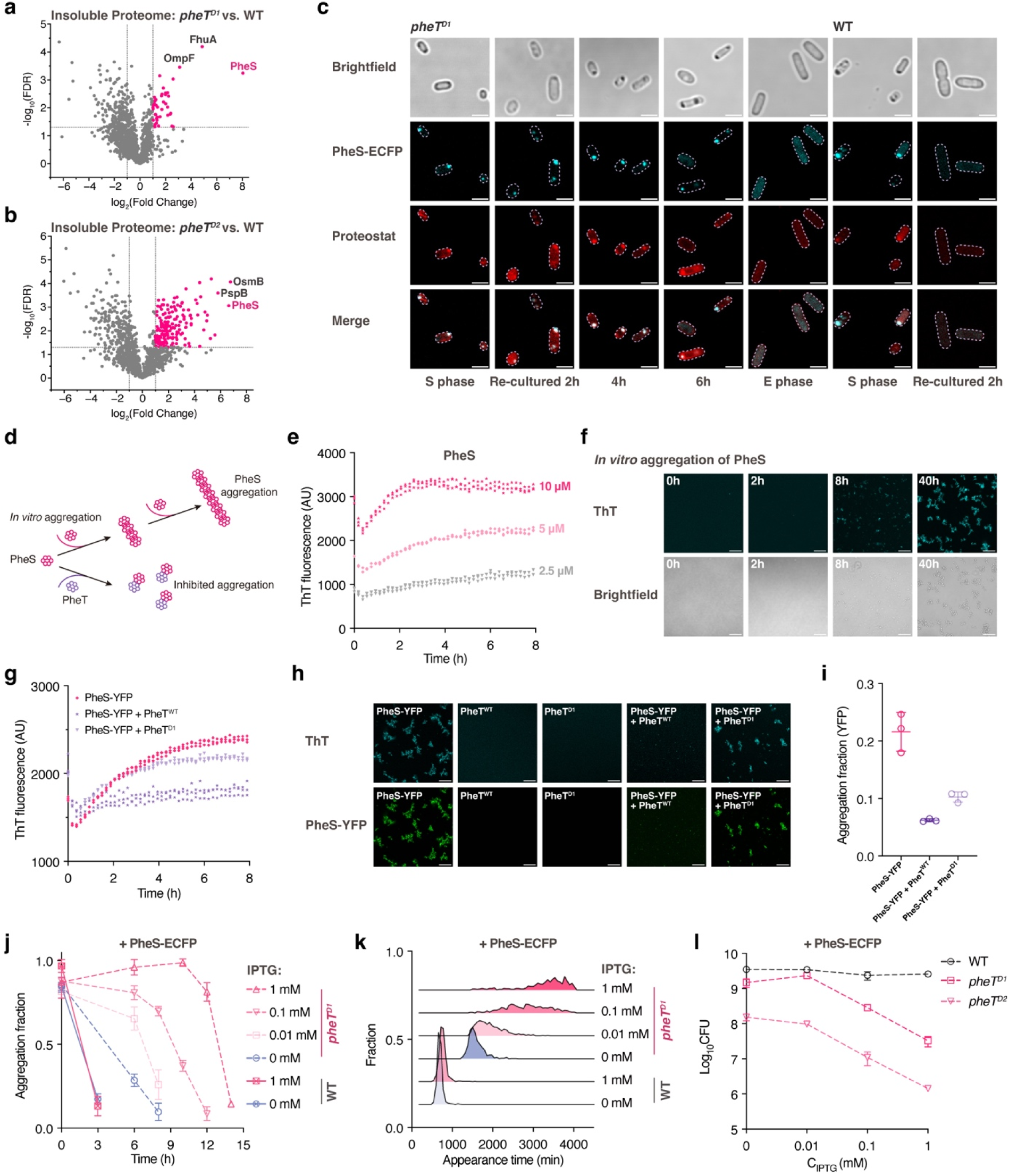
Increased PheS aggregation delays aggregate disassembly and extends lag time. **a**, **b**, Volcano plots showing MS analysis of the insoluble proteome in the *pheT^D1^*(**a**) and *pheT^D2^*(**b**) relative to WT at the stationary phase. Student’s *t*-test with the correction to multiple hypotheses by FDR-adjusted *p* < 0.05. n = 3 biological replicates. **c**, Representative brightfield and fluorescence images of WT and *pheT^D1^* cells expressing ECFP-tagged PheS and stained with Proteostat. Cells from the stationary phase were re-cultured in fresh LB medium and stained with Proteostat at the indicated time points. Scale bars, 2 μm. **d**, Schematic illustration of the PheS aggregation system *in vitro*. **e**, Fluorescence ThT signal over time for purified PheS at the indicated concentrations. Purified PheS was incubated in buffer containing 20[mM Tris at pH 7.4 and the corresponding NaCl concentrations of 38 mM, 19 mM, and 9.5 mM for 10 μM, 5 μM, and 2.5 μM PheS, respectively. Each point represents the normalized ThT fluorescence of one independent sample at each time point. n = 3 independent samples. **f**. Representative brightfield and ThT fluorescence images of purified PheS after the indicated times of incubation. PheS was incubated at 5 μM in buffer containing 20[mM Tris at pH 7.4 and 19 mM NaCl. Scale bars, 20 μm. **g**, Fluorescence ThT signal over time during co-incubation of PheS-YFP with PheT^WT^ or PheT^D1^. Each protein was used at 5 μM in buffer containing 20[mM Tris at pH 7.4 and 32 mM NaCl. Each point represents the normalized ThT fluorescence of one independent sample at each time point. n = 3 independent samples. **h**, Representative fluorescence images of the indicated samples after 48h incubation. PheS-YFP, PheT^WT^, and PheT^D1^ were each used at 5 μM. Scale bars, 20 μm. **i**, Quantitative sedimentation analysis of YFP fluorescence in the pellet fraction. Samples shown in (**g**) were incubated for 48h and then fractionated into supernatants and pellets by centrifugation at 15000g for 20 min. The fraction of YFP fluorescence recovered in the pellet was then quantified. Data are presented as mean ± SD (n = 3 independent samples). **j**, Quantification of the fraction of WT and *pheT^D1^* cells containing Proteostat-positive aggregates during recovery in fresh LB medium. Cells carrying an IPTG-inducible PheS-ECFP construct were grown for 48h at the indicated IPTG concentrations, and then transferred to fresh LB medium for recovery. Data are presented as mean ± SD (n = 4 biological replicates). **k**, Appearance time distribution for WT and *pheT^D1^* cells after induction of PheS-ECFP expression for 48h at the indicated IPTG concentrations. Sample sizes N = 1063, 694, 635, 1247, 1111, and 755, respectively. **l**, CFU of WT, *pheT^D1^*, and *pheT^D2^* cells after induction of PheS-ECFP expression for 48 h at the indicated IPTG concentrations. Data are presented as mean ± SD (n = 3 biological replicates).

To determine whether PheS can self-assemble into protein aggregates *in vitro*, we purified PheS and monitored its aggregation kinetics using thioflavin T (ThT) (Fig. 2d), a fluorescent probe whose signal increases upon binding to β-sheet-rich protein aggregates^25,26^. When incubated alone at room temperature, PheS exhibited a time-dependent increase in ThT signal (Fig. 2e and Extended Data Fig. 3d), accompanied by the progressive formation of ThT-positive aggregates under fluorescence microscopy (Fig. 2f). We observed that PheS aggregates formed over a broad range of conditions tested, with the extent of aggregate formation being further enhanced at higher PheS concentrations and lower salt concentrations (Fig. 2e and Extended Data Fig. 3d). By contrast, neither WT nor mutant PheT showed detectable ThT fluorescence increase or aggregate formation when incubated alone (Extended Data Fig. 3e, f). Thus, these observations suggest a previously unrecognized intrinsic propensity of PheS for self-aggregation.

We next tested whether PheT modulates PheS aggregation *in vitro*. PheS-YFP alone readily formed ThT-positive aggregates, whereas co-incubation with PheT^WT^ strongly suppressed aggregate formation, with less increase in ThT signal over time and no obvious aggregates detected microscopically (Fig. 2g, h). In contrast, the mutant PheT^D1^ retained only partial anti-aggregation activity compared to PheT^WT^ (Fig. 2g, h). Quantitative sedimentation analysis further revealed that PheT^WT^ markedly reduced PheS-YFP partitioning into the pellet, whereas PheT^D1^ was less effective (Fig. 2i). Together, these results reveal a chaperone-like activity of PheT in protecting PheS from aggregation.

To establish a causal relationship between PheS aggregation and dormancy, we examined how titrating PheS levels modulated aggregate dynamics and lag time. Bacteria carrying an IPTG-inducible PheS-ECFP system were grown for 48 hours in the stationary phase under graded concentrations of IPTG. Quantitative analysis of PheS-ECFP and Proteostat fluorescence showed that elevated PheS expression did not increase the Proteostat signal in WT cells (Extended Data Fig. 4a), paralleling the anti-aggregation activity of PheT^WT^ toward PheS. In contrast, with the induction of PheS, the *pheT* mutants exhibited a dose-dependent increase in insoluble aggregates (Extended Data Fig. 4a). Upon transfer to fresh medium, the elevated PheS aggregation burden dose-dependently delayed aggregate disassembly (Fig. 2j) and produced corresponding extensions of lag times (Fig. 2k and Extended Data Fig. 4b). This dormancy-promoting effect was independent of the canonical function of PheS in translation, as a PheS variant with abolished tRNA-binding sites^24^ comparably extended lag time (Extended Data Fig. 4c). At higher levels of PheS induction, the extended lag time progressed into a loss of culturability, characterized by a remarkable reduction in colony-forming units (CFUs) (Fig. 2l). This PheS-induced extension of lag time further enhanced ETP tolerance and compromised the cellular ability to cope with proteotoxic stress (Extended Data Fig. 4d, e). Thus, bacterial lag time can be fine-tuned by modulating the aggregation level of PheS.

### Protein aggregation sequesters the replication initiator DnaA to inhibit DNA synthesis

We further sought to elucidate the mechanism by which increased PheS aggregation prolongs dormancy. Previous studies reported that chaperones and proteases could facilitate protein disaggregation and cell resuscitation^4,12,27^. Nevertheless, we found that overexpressing chaperones (like DnaK, ClpB) failed to accelerate growth resumption, and overexpressing proteases (HslVU, Lon) unexpectedly further delayed dormancy exit in WT and two *pheT* mutants (Fig. 3a and Extended Data Fig. 5a). We observed that the protease component HslU formed aggregates colocalized with PheS-ECFP and showed synchronized delayed disaggregation in two *pheT* mutants upon nutrient replenishment (Extended Data Fig. 5b). It has been reported that proteases can degrade protein aggregates^28–30^. However, if so, their overexpression would be expected to accelerate, rather than delay, the exit from dormancy.

**Fig. 3.**
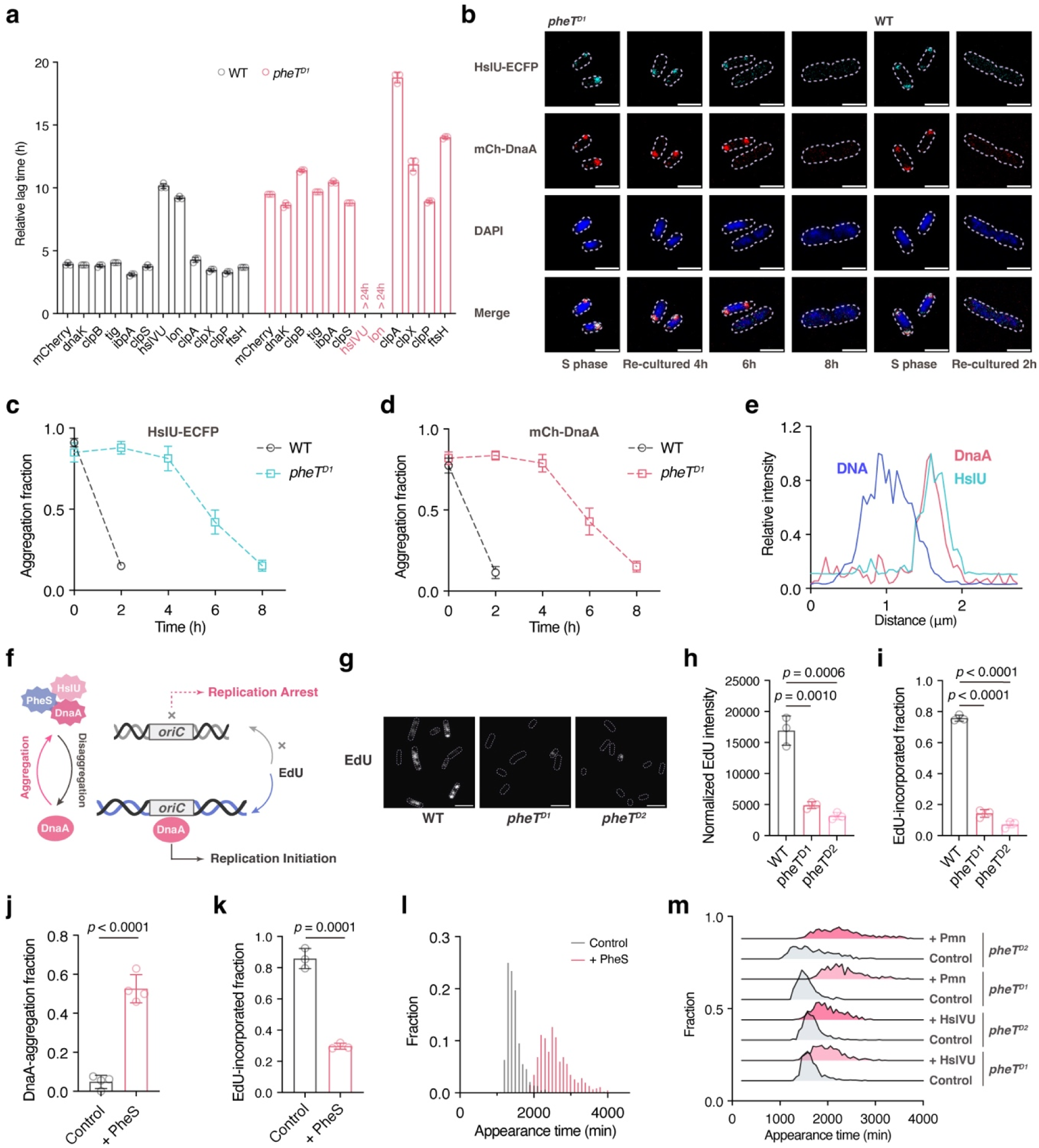
Sequestration of the replication initiator DnaA within aggregates inhibits DNA synthesis and extends lag time. **a**, Effect of overexpressing chaperones and proteases on dormancy exit of WT and *pheT^D1^* cells. Relative lag time was determined as the last time point at which ΔOD_600_ remained below 0.02. Data are presented as mean ± SD (n = 3 biological replicates). **b**, Representative fluorescence images of cells carrying chromosomally labeled HslU-ECFP and mCherry-DnaA stained with DAPI upon re-culturing. Cells from the stationary phase were re-cultured in fresh LB medium and sampled at the indicated time points. Scale bars, 2 μm. **c**, **d**, Quantification of the fraction of WT and *pheT^D1^* cells containing HslU-ECFP foci (**c**) or mCherry-DnaA foci (**d**) upon re-culturing. Data are presented as mean ± SD (n = 4 biological replicates). **e**, Representative relative intensity profiles of HslU-ECFP, mCherry-DnaA, and DNA (DAPI) in bacteria containing protein aggregates. Fluorescence intensity was measured along a line across the cell long axis. **f**, Schematic illustration of the replication initiation arrest through PheS-HslU-DnaA aggregation. **g**, Representative fluorescence images of WT, *pheT^D1^*, and *pheT^D2^* cells stained with Click-iT EdU Imaging Kit. Before staining, cells from the stationary phase were re-cultured in fresh LB medium with the addition of 30 μg/mL EdU for 60 minutes. Scale bars, 4 μm. **h**, **i**, Quantification of Normalized EdU intensity (**h**) and EdU-incorporated cell fraction (**i**) after Click-iT EdU staining in (**g**). Data are presented as mean ± SD (n = 3 biological replicates). *P* values were calculated using unpaired two-tailed t-tests. **j**, Quantification of the fraction of *pheT^D1^* cells containing DnaA aggregation after recovery in fresh LB medium for 7 hours with or without the induced expression of PheS. The IPTG concentration used was 1 mM. Data are presented as mean ± SD (n = 4 biological replicates). *P* values were calculated using unpaired two-tailed t-tests. **k**, Quantification of the EdU-incorporated cell fraction of the *pheT^D1^* cells with or without the induced expression of PheS. Cells were re-cultured in fresh LB medium for 6 hours, and 30 μg/mL EdU was added to label bacteria for 60 minutes. The IPTG concentration used was 1 mM. Data are presented as mean ± SD (n = 3 biological replicates). *P* values were calculated using unpaired two-tailed t-tests. **l**, Appearance time distribution for the *pheT^D1^* cells with or without the induced expression of PheS with 1 mM IPTG. Sample sizes N = 1164 and 468. **m**, Appearance time distribution for *pheT^D1^* and *pheT^D2^* cells during induced expression of HslVU-mCherry with 0.15 mM IPTG or treatment with 16 μg/mL puromycin. Sample sizes N = 673, 707, 622, 401, 810, 642, 678, and 402.

A recent study showed that depletion of the protease Lon abolished the extended lag time in a *metG* mutant^5^, supporting the function of proteases in promoting dormancy. Beyond their role in aggregate clearance, increasing evidence suggests that proteases also actively target and degrade native proteins involved in cell-cycle regulation^1,27,31^. In *Caulobacter crescentus*, proteases are activated by misfolded proteins to degrade the chromosomal replication initiator DnaA and induce cell-cycle arrest^1^. Given the conserved role of DnaA in replication initiation and its specificity as a protease substrate in *E. coli*^32,33^, we next explored whether protein aggregation might regulate cell-cycle progression.

We constructed chromosomal fluorescent fusions of HslU and DnaA and confirmed that this labelling did not perturb the lag-time distributions of WT or *pheT^D1^* mutant (Extended Data Fig. 5c). In stationary-phase WT and *pheT^D1^* cells, we observed that mCherry-DnaA formed aggregates that colocalized with HslU-ECFP (Fig. 3b). Upon nutrient replenishment, these aggregates rapidly dissolved in WT cells, whereas DnaA remained sequestered within aggregates for more than six hours in the *pheT^D1^* cells (Fig. 3b–d). Protein aggregates have been shown to localize to nucleoid-free regions owing to macromolecular crowding by the nucleoid^34,35^, so we examined the spatial relationship between DnaA and chromosomal DNA by DAPI staining. Remarkably, DnaA aggregates remained spatially separated from the nucleoid throughout the lag phase (Fig. 3b–e). To evaluate the effects of this segregation on DNA replication, newly synthesized DNA was monitored using incorporation of the thymidine analogue 5-ethynyl-2′-deoxyuridine (EdU) followed by Click-iT detection (Fig. 3f). Within 60 min of incubation, EdU incorporation was detected in approximately 80% of WT cells but remained rare in both *pheT* mutants (Fig. 3g–i), indicating suppressed DNA synthesis in dormant populations. These results support that persistent aggregates sequester DnaA away from the nucleoid to inhibit DNA replication during dormancy exit.

We therefore tested whether increasing proteostasis burden would further prolong DnaA sequestration and delay replication initiation. We found that elevated PheS expression further delayed the disassembly of DnaA aggregates (Fig. 3j), suppressed DNA synthesis (Fig. 3k), and extended lag time (Fig. 3l). Besides PheS, proteotoxic stress induced by elevated HslVU expression or puromycin produced similar lag-time extensions (Fig. 3m), suggesting that the proteostasis-replication axis could integrate distinct proteotoxic perturbations to tune bacterial dormancy. Moreover, combining these proteotoxic stresses synergistically promotes a dose-dependent reduction in colony-forming capacity (Extended Data Fig. 5d, e). These findings imply that the proteostasis decline could establish a continuum from extended lag times to an unculturable state by progressively restricting DnaA availability for replication initiation.

### Tunable DnaA co-aggregation quantitatively links proteostasis to replication initiation

We then explored whether reducing DnaA sequestration could facilitate replication initiation and dormancy exit. Previous studies implied that the position of fluorescent tags could affect DnaA localization and activity^36,37^, so we compared DnaA variants carrying mCherry at the amino terminus, carboxy terminus, or an internal site^36^. Notably, we found that the carboxy-terminal DnaA-mCherry variant showed substantially less aggregation than the other variants in stationary-phase *pheT^D1^* cells containing HslU-marked aggregates (Fig. 4a, b), which provides a more soluble variant to test the role of DnaA sequestration in dormancy exit.

**Fig. 4.**
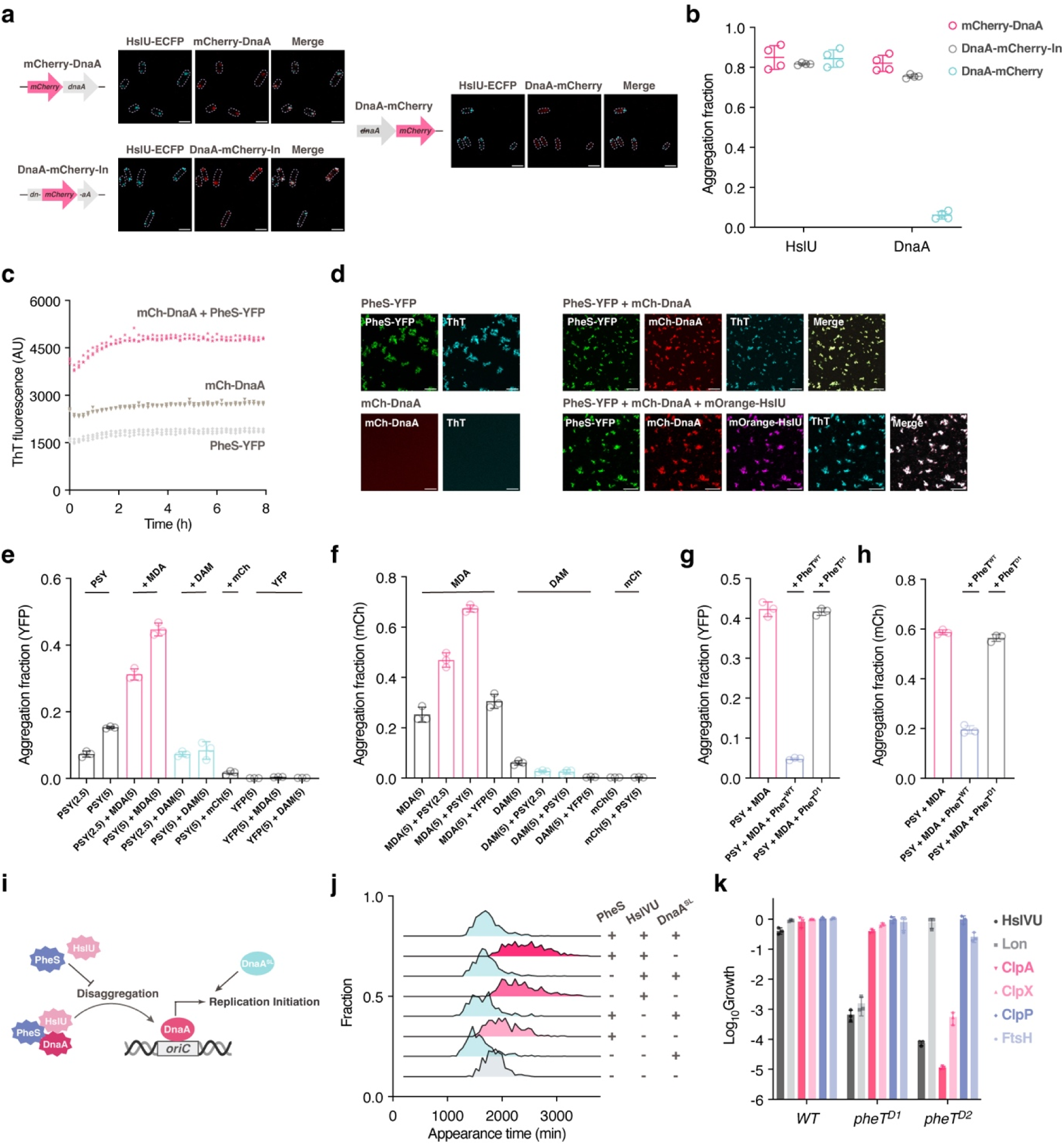
Tunable PheS-DnaA co-aggregation links aggregate burden to bacterial lag time. **a**, Representative fluorescence images of the *pheT^D1^* cells carrying chromosomal fluorescent fusions of HslU and DnaA at the stationary phase (48-hour cultures). mCherry-DnaA, DnaA with amino-terminus labelled mCherry; DnaA-mCherry-In, DnaA with internal-site labelled mCherry; DnaA-mCherry, DnaA with carboxy-terminus labelled mCherry. Scale bars, 2 μm. **b**, Quantification of the fraction of *pheT^D1^* cells containing PheS-ECFP aggregation and the indicated DnaA-variant aggregation at the stationary phase (48-hour cultures). Data are presented as mean ± SD (n = 4 biological replicates). **c**, Fluorescence ThT signal over time during incubation of the indicated protein samples. Each protein was used at 5 μM in buffer containing 20[mM Tris at pH 7.4 and 62 mM NaCl. Each point represents the normalized ThT fluorescence of one independent sample at each time point. n = 3 independent samples. **d**, Representative fluorescence images of the indicated samples after 48h incubation. PheS-YFP, mCherry-DnaA, and mOrange-HslU were each used at 5 μM in buffer containing 20[mM Tris at pH 7.4 and 62 mM NaCl. Scale bars, 20 μm. **e**, Quantitative sedimentation analysis of YFP fluorescence in the pellet fraction after 48h incubation. PheS-YFP (PSY) was incubated at a concentration of 2.5 μM or 5 μM alone or in the presence of mCherry-DnaA (MDA), DnaA-mCherry (DAM), or mCherry (mCh) at the indicated concentration in buffer containing 20[mM Tris at pH 7.4 and 62 mM NaCl. As controls, 5 μM YFP was incubated with 5 μM MDA or DAM. Data are presented as mean ± SD (n = 3 independent samples). **f**, Quantitative sedimentation analysis of mCherry fluorescence in the pellet fraction after 48h incubation. MDA, DAM, or mCh was incubated at a concentration of 5 μM alone or in the presence of PSY or YFP at the indicated concentration in buffer containing 20[mM Tris at pH 7.4 and 62 mM NaCl. Data are presented as mean ± SD (n = 3 independent samples). **g**, **h**, Quantitative sedimentation analysis of YFP fluorescence (**g**) and mCherry fluorescence (**h**) in the pellet fraction after 48h incubation. PSY and MDA were incubated with PheT^WT^ or PheT^D1^ at 5 μM in buffer containing 20[mM Tris at pH 7.4 and 62 mM NaCl. Data are presented as mean ± SD (n = 3 independent samples). **i**, Schematic illustration of promoting replication initiation with the soluble DnaA variant. **j**, Appearance time distribution for *pheT^D1^* cells during induced expression of PheS with 1 ng/mL anhydrotetracycline, HslVU-ECFP with 100 mM arabinose, or soluble DnaA-mCherry with 0.25 mM IPTG. Sample sizes N = 516, 562, 335, 309, 691, 491, 388, and 374, respectively. **k**, Relative colony-forming fraction of WT, *pheT^D1^*, and *pheT^D2^* cells during induced expression of the indicated proteases with 0.25 mM IPTG, normalized to IPTG-free plates. Data are presented as mean ± SD (n = 3 biological replicates).

To further understand the molecular basis of PheS-DnaA co-aggregation, we reconstituted their interaction *in vitro* (Fig. 4c, d). PheS-YFP alone readily formed ThT-positive aggregates, whereas mCherry-DnaA alone showed no detectable aggregation under the same conditions (Fig. 4d). Notably, co-incubation of PheS-YFP and mCherry-DnaA markedly increased ThT fluorescence and generated assemblies containing both proteins (Fig. 4c, d). Sedimentation assays further showed increased pelletable fractions of both PheS-YFP and mCherry-DnaA upon co-incubation, indicating a synergistic promoting effect (Fig. 4e, f). As controls, neither co-incubation of YFP with mCherry-DnaA nor of mCherry with PheS-YFP promotes co-aggregation (Fig. 4e, f and Extended Data Fig. 6a–d). Increasing PheS-YFP concentrations dose-dependently promoted the aggregation of mCherry-DnaA (Fig. 4f), and this co-aggregation was markedly inhibited by PheT^WT^, but not by the mutant PheT^D1^(Fig. 4g, h). These results indicate a quantitative link between aggregate burden and DnaA sequestration.

We then examined whether the reduced aggregation of DnaA-mCherry *in vivo* reflected a lower propensity for PheS-dependent co-aggregation. Co-incubation of PheS-YFP with DnaA-mCherry still increased ThT fluorescence and generated colocalized assemblies (Extended Data Fig. 6e, f). Unlike mCherry-DnaA, however, DnaA-mCherry showed no progressive increase in its pelletable fraction with increasing PheS-YFP concentrations and did not further promote PheS-YFP aggregation (Fig. 4e, f). Thus, DnaA-mCherry retains the ability to associate with PheS aggregates but is markedly less prone to synergistic co-aggregation, paralleling its reduced aggregation *in vivo*.

Importantly, we wondered whether the soluble DnaA variant was sufficient to bypass the aggregation-mediated dormancy (Fig. 4i). In line with previous reports^33^, overexpressing native DnaA perturbed replication initiation and caused pronounced growth defects in bacteria (Extended Data Fig. 7a). Strikingly, in contrast, we found that overexpression of the soluble DnaA variant accelerated the dormancy exit of the *pheT^D1^* cells and completely abolished the extended lag times induced by elevated PheS or HslVU levels (Fig. 4j and Extended Data Fig. 7a). A carboxy-terminal fusion of DnaA to the solubility-enhancing maltose-binding protein (MBP) produced similar phenotypes (Extended Data Fig. 7b, c). Therefore, these rescue experiments support DnaA sequestration as a downstream mechanism through which aggregate burden converges to delay dormancy exit.

Taken together, these findings establish PheS-dependent DnaA co-aggregation as a tunable mechanism that regulates DnaA availability for replication initiation, whereas increasing DnaA solubility bypasses this sequestration and accelerates growth resumption.

### A branched proteostasis-replication network underlies dormancy control

We further investigated whether the PheS-DnaA co-aggregation could recruit additional aggregate-associated proteins. Although mOrange-HslU alone did not form obvious aggregates, co-incubation with PheS-YFP generated assemblies containing both proteins (Extended Data Fig. 7d). Among the conditions tested, co-incubation of PheS-YFP, mCherry-DnaA, and mOrange-HslU produced the largest increase in ThT fluorescence and generated co-aggregates containing all three proteins (Fig. 4d and Extended Data Fig. 7e). These findings indicate that proteins with distinct cellular functions can be incorporated into shared co-aggregated assemblies to regulate bacterial dormancy.

Nevertheless, we noticed that overexpressing soluble DnaA did not accelerate dormancy exit in the *pheT^D2^* mutant (Extended Data Fig. 7f), implying additional proteostasis-sensitive targets contribute to its extended lag time. As *E. coli* encodes multiple proteases, including HslVU, Lon, ClpP in complex with ClpX or ClpA, and FtsH^27^, we systematically examined how these proteases affected dormancy exit in different mutants. We found that, while both *pheT* mutants were sensitive to elevated HslVU, they showed distinct responses to other proteases: the *pheT^D1^* cells were more sensitive to Lon, whereas the *pheT^D2^* cells exhibited greater sensitivity to ClpA and ClpX (Fig. 4k). These results imply that bacteria may rely on multiple, partly redundant proteostasis pathways to coordinate stress responses with DNA replication.

The cysteine tRNA mutant *cysT^D1^*exhibited similarly compromised proteostasis capacity as shown before (Fig. 1a, c, h). Further proteomic analysis of the insoluble fractions revealed mutation-specific aggregate composition across the *pheT^D1^*, *pheT^D2^*, and *cysT^D1^* cells (Extended Data Fig. 8a, b). Despite this heterogeneity, the ferrichrome transporter FhuA^38^ showed consistently elevated aggregation across all three mutants (Extended Data Fig. 8a, b). We demonstrated that inducible expression of FhuA alone was sufficient to prolong dormancy in all these mutants (Extended Data Fig. 8c, d). This effect was independent of its canonical function in ferrichrome transport, as a transport-defective FhuA Δ236-248 variant^39^ produced a comparable lag extension (Extended Data Fig. 8e). Elevated expression of protease HslVU, Lon, or ClpA also further delayed dormancy exit in the *cysT^D1^*cells (Extended Data Fig. 8f).

Thus, these results imply a broader, branched proteostasis-replication network in which diverse proteotoxic stresses converge to tune the switch between dormancy and proliferation.

### A scaling relationship between lag extension and proteostasis decline

We then tested whether the proteostasis-governed dormancy extends from genetic perturbations to environmentally induced proteotoxic stress. We observed that long-lag WT cells induced by heat shock or puromycin retained HslU and DnaA within aggregates throughout dormancy (Fig. 5a), indicating the generality of the proteostasis-replication axis across environmental conditions. Exposure to 55°C for 1 h delayed aggregate disassembly and prolonged lag time, whereas treatment at 49°C for 1 h, despite being sufficient to induce aggregates in fast-growing cells (Extended Data Fig. 2b), had little effect on either aggregate disassembly or lag duration (Fig. 5b, c). These results suggest that the bacterial protein quality-control network can buffer low levels of proteotoxic stress to support timely regrowth, and lag time is extended only when cumulative stress overwhelms cellular proteostasis capacity to manage aggregates. Supporting this view, at concentrations below the minimum inhibitory concentration (MIC), puromycin permitted bacterial growth but extended lag times in a dose-dependent manner (Extended Data Fig. 9a). Following treatment with puromycin above the MIC, lag time after drug removal increased progressively with previous exposure duration (Fig. 5b–d), suggesting a cellular memory of accumulated proteotoxic burden. Notably, increasing soluble DnaA accelerated recovery from puromycin-induced dormancy (Extended Data Fig. 9b), further supporting DnaA as a common gatekeeper of dormancy exit.

**Fig. 5.**
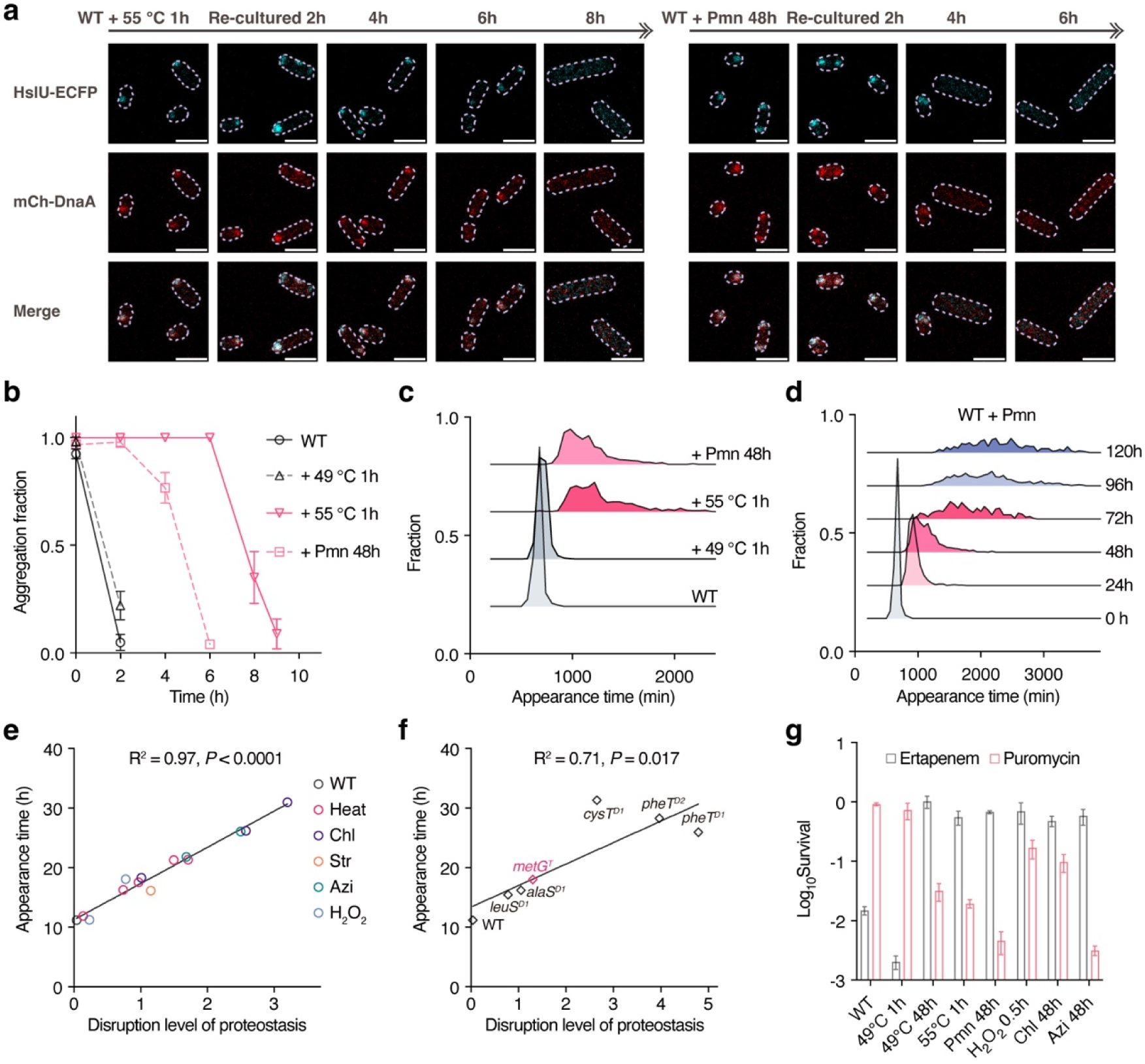
Lag time scales with proteostasis decline across different backgrounds. **a**, Representative fluorescence images of cells carrying chromosomally labeled HslU-ECFP and mCherry-DnaA after exposure to the indicated proteotoxic stress and re-culturing with fresh LB medium. Stationary-phase WT cells were treated with 256 μg/mL puromycin for 48h or at 55 °C for 1h in fresh LB medium. Upon stress removal, bacteria were re-cultured with fresh LB medium and sampled at the indicated time points. Scale bars, 2 μm. **b**, Quantification of the fraction of stress-treated WT cells containing Proteostat-positive aggregates upon re-culturing. WT cells were treated with the indicated stress, and upon stress removal, bacteria were re-cultured with fresh LB medium and stained with Proteostat at the indicated time points. Data are presented as mean ± SD (n = 4 biological replicates). **c**, Appearance time distribution for WT after treatment with the indicated stress. Sample sizes N = 989, 736, 442, and 733, respectively. **d**, Appearance time distribution for WT after exposure to 256 μg/mL puromycin for the indicated durations. Sample sizes N = 989, 683, 733, 288, 653, and 369, respectively. **e**, Mean appearance time plotted against the disruption level of proteostasis after treatment with the indicated stress. Stationary-phase WT cells were diluted in fresh LB medium supplemented with the indicated stress conditions. From left to right: Heat: 49 °C for 1h, 24h, or 48h; 55 °C for 0.5h or 1h; Chloramphenicol (Chl): 500 μg/mL for 48h, 72h, 96h; Streptomycin (Str): 50 μg/mL for 2h; Azithromycin (Azi): 100 μg/mL for 24h, 48h; H_2_O_2_: 5 mM or 10 mM for 0.5h. The disruption level of proteostasis is quantified as the log_10_ ratio of CFU on antibiotic-free versus 64 μg/mL puromycin plates. Each point represents the mean of n = 3 biological replicates for each stress condition. The solid line denotes linear regression (R^2^ = 0.97, *P* < 0.0001). **f**, Mean appearance time plotted against the disruption level of proteostasis for the identified dormant mutants. Each point represents the mean of n = 3 biological replicates for each mutant. The disruption level of proteostasis is quantified as the log_10_ ratio of CFU on antibiotic-free versus 64 μg/mL puromycin plates. The solid line denotes linear regression (R^2^ = 0.71, *P* = 0.017). **g**, Relative colony-forming fraction of stress-treated WT cells on plates containing 64 μg/mL puromycin, normalized to drug-free plates, together with survival fraction after 4h of 1 µg/mL ertapenem treatment. Data are presented as mean ± SD (n = 3 biological replicates).

To compare the extent of proteostasis disruption across different environmental conditions, we attempted to use puromycin susceptibility, rather than aggregate abundance, as a functional indicator of proteostasis disruption based on the following observations. First, we have shown that puromycin can promote protein aggregation, delay aggregate disassembly, and extend lag time in a dose-dependent manner (Fig. 5a–d). Accordingly, progressive disruption of proteostasis would be reflected in a gradual decrease in the ability to cope with puromycin-induced aggregation. Second, dormant mutants contained a similar aggregate amount relative to WT cells (Extended Data Fig. 1g), but exhibited markedly increased puromycin sensitivity (Fig. 1h), indicating that puromycin sensitivity reflects proteostasis disruption more accurately than the aggregate abundance. Third, we confirmed that treatment at 49 °C for 1 hour failed to delay aggregate disassembly, consistent with its unaffected sensitivity to puromycin, while the 55 °C treatment both slowed aggregate disassembly and increased puromycin sensitivity (Fig. 5b, c and Extended Data Fig. 9c). Thus, puromycin sensitivity provides a functional readout of the proteostasis capacity in bacteria.

Using this measure, we quantified proteostasis disruption across diverse environmental perturbations as the log_10_ ratio of CFU recovered on antibiotic-free medium to that on puromycin-containing medium. Strikingly, we observed that lag duration increased linearly with the extent of proteostasis disruption, regardless of stress type (Fig. 5e). Prolonged exposure to azithromycin or chloramphenicol similarly increased puromycin susceptibility and extended lag time, indicating that this relationship was not restricted to canonical proteotoxic treatments (Fig. 5e). The quantitative relationship also extended to genetically dormant backgrounds, including the previously described long-lag mutant *metG^T^*^3^ (Fig. 5f). This scaling relationship suggests a universal trade-off between dormancy and proteostasis. Similar to dormant mutants, stress-induced dormancy also conferred increased ertapenem tolerance while causing collateral sensitivity to additional proteotoxic challenges (Fig. 5g). From a clinical perspective, the disrupted proteostasis could represent an Achilles’ heel of dormant bacteria, defining a shared vulnerability of these antibiotic persisters.

### Restoring proteostasis accelerates dormancy exit

Once antibiotic stress is removed, rapid resumption of growth should confer a significant fitness advantage. We therefore performed experimental evolution to identify genetic factors that accelerate dormant exit of the *pheT* mutants. Serial passaging of the *pheT* mutants in antibiotic-free LB medium rapidly selected populations with shortened lag times within seven cycles (Extended Data Fig. 10a, b). Reverse mutants were subsequently isolated and analysed by WGS (Extended Data Fig. 10c and Supplementary Table 2). Intriguingly, while these reverse mutations mapped to diverse gene families, improved proteostasis emerges as a convergent signature among reverse mutants, characterized by accelerated aggregate disassembly and reduced susceptibility to proteotoxic stress (Extended Data Fig. 10d, e). These results further support proteostasis as a major determinant of dormancy exit.

As delayed aggregate disassembly extends lag time, we reasoned that promoting aggregate disassembly could accelerate dormancy exit. Increasing evidence indicates that reactive oxygen species (ROS), particularly hydrogen peroxide, can form spontaneously at aggregate interfaces and establish a self-reinforcing feedback loop that promotes aggregation^40,41^. Using the H_2_O_2_-specific fluorogenic probe Peroxy Orange 1 (PO-1)^42,43^, we detected enrichment of H_2_O_2_ within aggregates compared to the cytoplasm (Extended Data Fig. 11a, b). Notably, the *pheT^D1^* cells maintained higher H_2_O_2_ levels than WT cells throughout dormancy (Extended Data Fig. 11a, b). Consistently, the *pheT^D1^* cells exhibited elevated sensitivity to Fe^2+^ treatment, which reacts with H_2_O_2_ via the Fenton reaction to generate more hydroxyl radicals^44^ (Extended Data Fig. 11c).

To determine whether elevated H_2_O_2_ contributes to delayed aggregate disassembly, we supplemented the medium with catalase to scavenge H_2_O ^45^ and monitored disaggregation dynamics. Remarkably, catalase treatment significantly reduced intracellular H_2_O_2_ levels in the *pheT^D1^* cells (Extended Data Fig. 11a, b) and accelerated aggregate disassembly (Extended Data Fig. 11d). Accordingly, the addition of catalase directly facilitated bacterial exit from dormancy (Extended Data Fig. 11e). Similar promoting effects were obtained with the supplement of thiourea, another well-characterized hydroxyl radical scavenger^46^ (Extended Data Fig. 11e). This effect was not restricted to genetically induced dormancy, as catalase also accelerated recovery from puromycin-induced dormancy (Extended Data Fig. 11f). Together, these findings show that scavenging aggregate-associated hydroxyl radicals promotes proteostasis recovery and accelerates dormancy exit.

### A universal coupling of proteostasis and dormancy across bacterial species

To test whether the co-aggregation of PheS and DnaA is specific to *E. coli* or extends to other bacterial species, we purified PheS and DnaA from *E. coli*, *Klebsiella pneumoniae*, and *Bacillus subtilis*. Sequence analysis revealed high similarity between the *E. coli* and *K. pneumoniae* proteins (~90%), whereas the corresponding *B. subtilis* proteins were substantially more divergent (~40%) (Fig. 6a). Despite this sequence divergence, the proteins exhibited similar aggregation behaviour: PheS-YFP readily formed ThT-positive aggregates when incubated alone, whereas mCherry-DnaA did not show obvious aggregated structures (Extended Data Fig. 12a). As observed for the *E. coli* proteins, co-incubation of PheS-YFP and mCherry-DnaA from either *K. pneumoniae* or *B. subtilis* increased ThT fluorescence, generated colocalized ThT-positive assemblies, and increased partitioning of both proteins into the pellet fraction (Fig. 6b–d and Extended Data Fig. 12b–e), indicating that PheS-DnaA co-aggregation is conserved across these species.

**Fig. 6.**
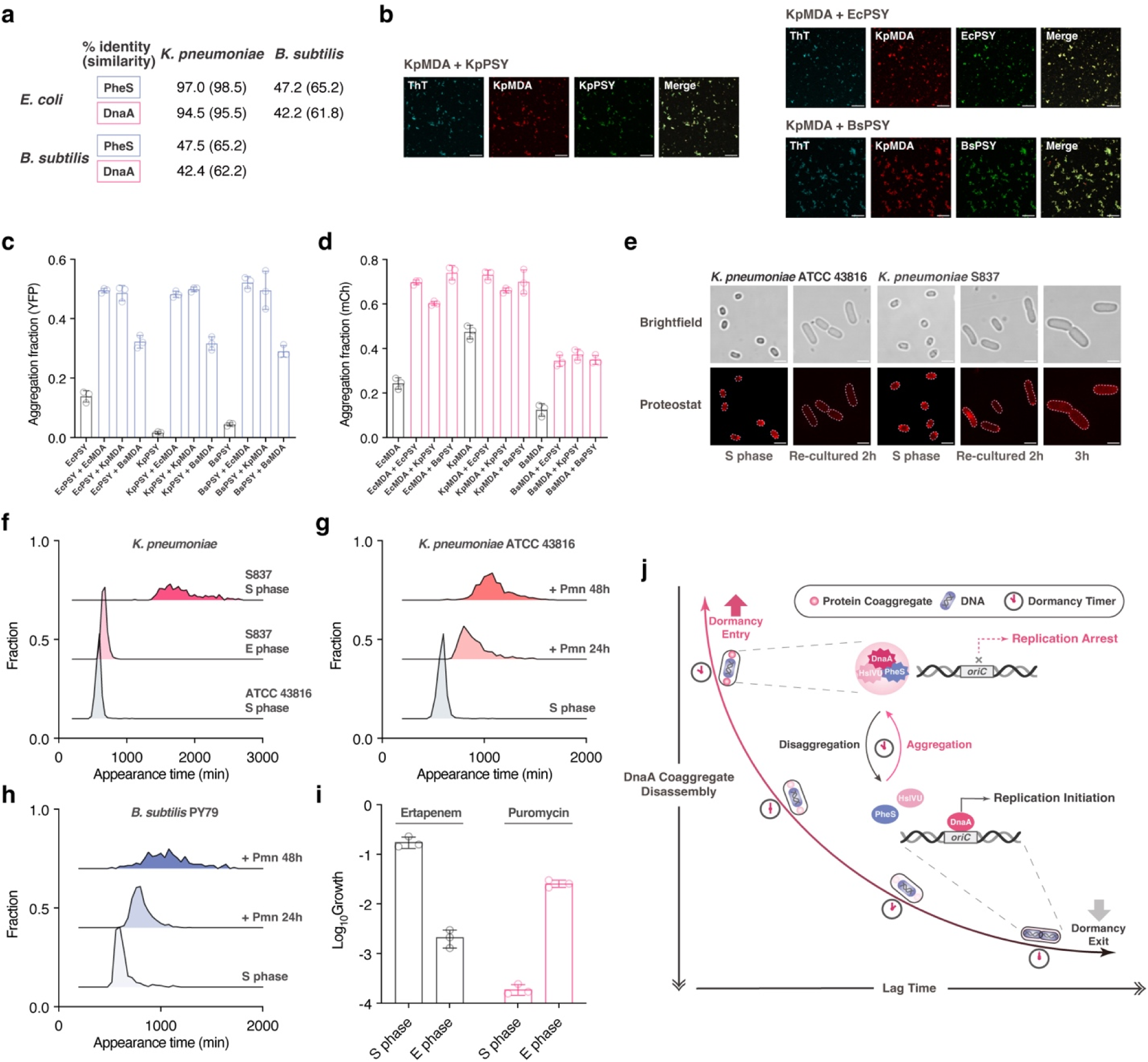
Conserved PheS-DnaA co-aggregation and proteostasis-dormancy coupling across bacterial species. **a**, Sequence identity and similarity (in brackets) of PheS from *E. coli*, *K. pneumoniae*, and *B. subtilis*, together with the corresponding values for DnaA from the three species. **b**, Representative fluorescence images of mCherry-DnaA and PheS-YFP from the indicated species after 48h co-incubation. Each protein was used at 5 μM in buffer containing 20[mM Tris at pH 7.4 and 62 mM NaCl. Scale bars, 20 μm. **c**, Quantitative sedimentation analysis of YFP fluorescence in the pellet fraction after 48h incubation. PheS-YFP (PSY) from *E. coli*, *K. pneumoniae*, or *B. subtilis* was incubated at 5 μM either alone or together with 5 μM mCherry-DnaA (MDA) from the indicated species in buffer containing 20[mM Tris at pH 7.4 and 62 mM NaCl. Data are presented as mean ± SD (n = 3 independent samples). **d**, Quantitative sedimentation analysis of mCherry fluorescence in the pellet fraction after 48h incubation. mCherry-DnaA (MDA) from *E. coli*, *K. pneumoniae*, or *B. subtilis* was incubated at 5 μM either alone or together with 5 μM PheS-YFP (PSY) from the indicated species in buffer containing 20[mM Tris at pH 7.4 and 62 mM NaCl. Data are presented as mean ± SD (n = 3 independent samples). **e**, Representative brightfield and fluorescence images of *K. pneumoniae* ATCC 43816 and clinical *K. pneumoniae* S837 stained with Proteostat. Bacteria from the stationary phase were re-cultured in fresh LB medium and sampled at the indicated time points. Scale bars, 2 μm. **f**, Appearance time distribution for clinical isolate *K. pneumoniae* S837 at the stationary phase or exponential phase, together with laboratory strain *K. pneumoniae* ATCC 43816 at the stationary phase. Sample sizes N = 1239, 802, and 293, respectively. **g**, **h**, Appearance time distribution for *K. pneumoniae* ATCC 43816 (**g**) and *B. subtilis* PY79 (**h**) after exposure to 256 μg/mL puromycin for the indicated durations. Sample sizes N = 1239, 1013, and 625 in (**g**). Sample sizes N = 438, 1287, and 181 in (**h**). **i**, Relative colony-forming fraction of stationary-phase and exponential-phase *K. pneumoniae* S837 on plates containing 64 μg/mL puromycin, normalized to drug-free plates, together with survival fraction after 3h of 1 µg/mL ertapenem treatment. Data are presented as mean ± SD (n = 3 biological replicates). **j**, A proteostasis clock controls bacterial lag time by timing replication initiation. Upon entry into dormancy, protein aggregates PheS and HslVU sequester the chromosome replication initiator DnaA away from the bacterial nucleoid to prevent replication initiation. After nutrient replenishment, replication initiation and DNA synthesis remain inhibited until DnaA aggregates are disassembled. The disassembly rate of DnaA aggregates therefore determines the duration of bacterial dormancy.

We next examined whether PheS and DnaA from different species could also co-aggregate. Strikingly, all heterologous PheS-DnaA combinations tested also produced time-dependent increases in ThT fluorescence, colocalized aggregates, and increased pelletable fraction (Fig. 6b–d and Extended Data Fig. 12b–e). Thus, synergistic aggregation of PheS and DnaA is not limited to proteins from the same species but extends across species boundaries, suggesting that the underlying interaction is broadly conserved during evolution.

Finally, Proteostat staining revealed the formation of protein aggregates in stationary-phase *K. pneumoniae* cells, and these aggregates disassembled more slowly after nutrient replenishment in the long-lag clinical isolate *K. pneumoniae* S837 than in the laboratory strain *K. pneumoniae* ATCC 43816 (Fig. 6e, f). Furthermore, we observed that puromycin-induced proteotoxic stress was sufficient to extend lag times in both *K. pneumoniae* and *B. subtilis* (Fig. 6g, h). In the clinical *K. pneumoniae* isolate S837, extended lag time was accompanied by increased ETP tolerance but greater susceptibility to puromycin (Fig. 6i), highlighting disrupted proteostasis as a shared vulnerability of antibiotic-tolerant populations.

Together, these findings reveal a conserved coupling between proteostasis and dormancy across different bacterial species and identify the proteostasis-replication network as a potential therapeutic target against antibiotic persisters.

## DISCUSSION

The accumulation of protein aggregates is typically linked to cytotoxicity and aging-related diseases in eukaryotes^19–21^. The disassembly of such aggregates is often viewed as too slow to be relevant on physiological timescales^47^. However, our work proposes that the kinetics of aggregate disassembly can be repurposed as a tunable timing device in bacteria for fine-tuning lag time to match the duration of stress (Fig. 6j). Conceptually, this proteostasis-governed timing device and canonical circadian clocks driven by transcription-translation feedback loops^48^ represent two distinct evolutionary solutions to the problem of temporal control. Circadian clocks provide a stable, cell-autonomous mechanism to synchronize internal physiology with the 24-hour light-dark cycle, whereas the proteostasis clock integrates the history of proteotoxic stress and encodes the timescale from disrupted proteostasis to a replication-capable state. From an evolutionary perspective, while dedicated oscillators are favoured in environments with regular cycles, a highly tunable timer based on the proteostasis network offers a more economical and flexible solution for bacteria adapting to fluctuating and unpredictable environments.

While protein aggregation has been implicated as a bacterial strategy to facilitate dormancy and antibiotic tolerance^49^, the mechanistic link between aggregates and cell-cycle progression has remained elusive. Our results suggest that protein aggregation is not simply a passive process of deposition but actively regulates the spatiotemporal dynamics of the replication initiator DnaA, thereby gating the transition from dormancy to replication. Guided by this, bacterial lag time can be precisely tuned by genetically or chemically targeting components of the proteostasis-replication axis. While DnaA emerges here as a critical client, other cell-cycle regulators are likely to participate in this timing mechanism, whose full regulatory architecture has yet to be resolved.

Our work shows that protein aggregation *per se* represents cellular responses to proteotoxic stresses, rather than a definitive indicator of dormancy. Instead, lag time distribution is determined by the disassembly rate of aggregates, a kinetic parameter that reflects the cellular proteostasis capacity. In bacteria, proteostasis is maintained by an intricate network that coordinates protein synthesis, folding, trafficking, and degradation^50^. This systemic nature may explain why long-lag phenotypes are associated with such diverse genetic mutations and environmental factors. These perturbations may represent distinct ways that convergently shift the global proteostasis network away from a replication-competent state. In this view, lag time can be understood as a manifestation of how far the proteostasis network has been driven away from the replication state.

Across *E. coli* and clinical *K. pneumoniae* isolates, extension of lag time confers ertapenem tolerance while simultaneously causing collateral sensitivity to proteotoxic stress, revealing a universal trade-off between dormancy and proteostasis. This trade-off highlights proteostasis pathways as promising targets for the clinical elimination of antibiotic persisters, providing two directions for combating dormant bacteria. One approach is to promote protein disaggregation to accelerate dormancy exit, thereby resensitizing bacteria to antibiotics that target actively proliferating cells. An alternative strategy is to exacerbate proteotoxic stress, which produces lag extension and ultimately drives cells toward a loss of culturability.

Beyond bacteria, protein aggregates have been observed in diverse dormant systems^51^, including yeast^52^, stem cells^17^, and oocytes^53^. The evolutionary conservation of proteostasis and DNA replication raises the possibility that regulated proteostasis may contribute more broadly to the timing of cellular dormancy. Heterotypic protein co-aggregation has been studied most extensively in neurodegenerative disease^54^, where one amyloid-like protein can promote the aggregation of another, as illustrated by the synergistic fibrillization of tau and α-synuclein^55^. The observation that a similar principle operates in bacterial dormancy is therefore unexpected and intriguing. One possibility is that these amyloid-like proteins could integrate prior cellular experience into history-dependent aggregation states and thereby encode a form of protein memory^56^. In this view, similar logic may underlie both pathological progression in neurodegeneration and dormancy timekeeping in bacteria.

## METHODS

### Bacterial strains, plasmids, and growth conditions

All strains used in this study are indicated in Supplementary Table 3, plasmids in Supplementary Table 4, and primers in Supplementary Table 5. *K. pneumoniae, B. subtilis, E. coli* KLY, and derivative strains were grown at 37 °C in Luria-Bertani (LB) broth (Miller formulation; 10 g/L NaCl, 10 g/L tryptone, and 5 g/L yeast extract) with 220 rpm shaking or on solid LB medium prepared by supplementing the broth with 15 g/L agar, unless otherwise stated.

Polymerase chain reaction (PCR) amplification was performed using PrimeSTAR Max DNA Polymerase (Takara) according to the manufacturer’s instructions. For plasmid construction, the gene PCR product was amplified from *E. coli* KLY and inserted into the pET28a/pET28a-mCherry backbone using the ClonExpress II One Step Cloning Kit (Vazyme). Cells were plated on LB plates containing kanamycin (100 μg/mL) for selection. Positive clones were validated by Sanger sequencing.

To label HslU and DnaA with fluorescent proteins on the chromosome, we used a two-step λ Red recombineering. In the first step, the *kan::sacB* cassette was amplified and transformed into the electrocompetent *E. coli* KLY cells containing IPTG-induced Red proteins (pSC101-RED-tet). Cells were plated on LB plates containing kanamycin (100 μg/mL) for selection. In the second step, the *hslU/dnaA-ECFP/mCherry* translational fusion fragment was introduced by electroporation into cells carrying the *kan::sacB* cassette at the targeting site, in which Red proteins had been induced with IPTG. Recombinants were counter-selected on sucrose plates and validated by Sanger sequencing.

To restore the WT alleles in the *pheT* mutants, P1 phage (ATCC 25404-B1) was used to perform general transduction^57^, and primer design was done using Duplication-Insertion recombineering^58^.

### Evolutionary protocol

For antibiotic evolution, an overnight culture was diluted 1:100 into 1 mL LB medium supplemented with 1 μg/mL ertapenem and incubated for 4 or 12 hours alternately at 37 °C with shaking. After killing, the cultures were washed twice with 0.9% NaCl to remove antibiotics, resuspended in 1 mL fresh LB medium, and then grown overnight at 37 °C with shaking. For reverse evolution, overnight cultures of the dormant mutants were diluted 1:100 into 1 mL antibiotic-free LB medium and serially passaged overnight at 37 °C with shaking.

### ScanLag analysis

Bacteria were serially diluted to approximately 200 CFU per plate and plated on solid LB medium. The plates were then placed in a ScanLag setup^14^ at 37 °C. This setup automatically monitors colony growth every 20 minutes and extracts the distribution of appearance times. The colony detection threshold size is set at 100 pixels.

### Whole genome sequencing

Genomic DNA was extracted using a TIANamp Bacteria DNA Kit (TIANGEN; Beijing, China). Sequencing was performed by Sinobiocore using Illumina HiSeq with 150 bp paired-end at an average coverage of 100-200. Single nucleotide polymorphisms (SNPs) and indels were identified by bwa version 0.7.18.

### Cell staining and microscopy

For Proteostat staining, bacteria were processed following the manufacturer’s instructions (Enzo Life Sciences; enz-51035-K100): Briefly, bacteria were fixed with 4% PFA for 30 minutes at room temperature and permeabilized in Assay Buffer with 0.5% Triton X-100 for 30 minutes on ice. Cells were then washed twice with PBS and stained with Proteostat dye for 30 minutes at room temperature. The stained cells were washed twice with PBS before imaging.

For DAPI staining, fixed cells were stained with DAPI (MCE, HY-D0814) at a concentration of 1 μg/mL in PBS for 10 minutes at room temperature. For Peroxy Orange 1 (PO-1) staining, bacteria were stained with PO-1 (MCE, HY-103469) at a concentration of 10 μM in PBS for 1 hour at 37 °C. The stained cells were washed twice with PBS before imaging.

Brightfield and fluorescence images were captured using a FV3000 confocal microscope (Olympus) with a 100 × objective. Appropriate filter sets were chosen for each fluorophore based on their excitation and emission spectra. All images were analyzed using Fiji.

### Insoluble protein isolation

The isolation of insoluble proteins was modified from Tomoyasu^16^. Briefly, cells (1 mL, 2 × 10^9^ bacteria) were collected and incubated in 40 μL buffer A (10 mM potassium phosphate buffer, pH 6.5, 1 mM EDTA, 20% (w/v) sucrose, 1 mg/mL lysozyme) for 30 minutes on ice. Cells were then mixed with 360 μL buffer B (10 mM potassium phosphate buffer, pH 6.5, 1 mM EDTA) and lysed by sonication while cooling. Intact cells were removed by centrifugation (2000 g, 20 minutes, 4 °C). The insoluble cell fraction was isolated by centrifugation (15000 g, 20 minutes, 4 °C). The pellet fractions were resuspended in 400 μL buffer B with 2% NP40 to dissolve membrane proteins and isolated by centrifugation (15000 g, 30 minutes, 4 °C). NP40-insoluble pellets were washed twice with 400 μL buffer B and resuspended in 100 μL buffer B by brief sonication. The concentration of insoluble protein was determined using the BCA protein assay kit (Beyotime).

### Growth curves

Bacteria were diluted to 2 × 10^5^ cells/mL in a 96-well plate of LB medium. Growth measurements were performed by incubating the plate in a SPARK multimode microplate reader (Tecan) at 37 °C with shaking. OD_600_ was measured every 10 minutes.

### Killing assays

For ertapenem and H_2_O_2_ killing, bacteria were diluted 1:100 into LB medium supplemented with 1 µg/mL ertapenem or 10 mM H_2_O_2_ and incubated for the indicated time at 37 °C with shaking. For heat killing, bacteria were diluted 1:100 into LB medium and incubated for 30 minutes at 55 °C. The CFUs were evaluated by plating before and after stress exposure.

### Culturability tests

Tenfold dilutions of bacteria were prepared in PBS and subsequently plated on solid LB medium supplemented with the indicated concentrations of puromycin or IPTG. The CFUs were calculated after 72 hours of incubation at 37 °C.

### EdU-Click labelling of newly replicated DNA

Newly synthesized DNA was labeled using the Click-iT EdU Imaging Kit (Invitrogen C10340) according to the manufacturer’s instructions. Specifically, bacteria grown in LB medium were incubated with 30 μg/mL EdU for 60 minutes. Cells were fixed with 4% PFA for 15 minutes and permeabilized in PBS with 0.5% Triton X-100 for 15 minutes at room temperature. Cells were then washed with PBS and incubated in 100 μL of Click-iT reaction cocktail for 30 minutes at room temperature, protected from light. The stained cells were washed twice with PBS before imaging.

### Quantitative proteomics

The total protein of bacteria is extracted using the Bacterial Protein Extraction Kit (Sangon). The insoluble protein is isolated as previously mentioned. The proteins were separated by SDS-PAGE, respectively. The gel bands of interest were excised from the gel, reduced with 5 mM of DTT, and alkylated with 11 mM iodoacetamide, which was followed by in-gel digestion with sequencing grade modified trypsin at 37 °C overnight. The peptides were extracted twice with 0.1% trifluoroacetic acid in 50% acetonitrile aqueous solution for 30 min and then dried in a speedvac. Peptides were redissolved in 20 μL 0.1% trifluoroacetic acid, and 6 μL of extracted peptides were analyzed by Thermo Scientific Orbitrap Exploris 480.

For LC-MS/MS analysis, the peptides were separated by an 85-min gradient elution at a flow rate of 0.30 µl/min with a Thermo-Dionex Ultimate 3000 HPLC system, which was directly interfaced with a Thermo Scientific Orbitrap Exploris 480 mass spectrometer. The analytical column was a home-made fused silica capillary column (75 µm ID, 350 mm length) packed with C-18 resin (1.9 µm, Dr.Maisch GmbH). Mobile phase consisted of 0.1% formic acid, and mobile phase B consisted of 80% acetonitrile and 0.1% formic acid. Orbitrap Exploris 480 mass spectrometer was operated in the data-dependent acquisition mode using Xcalibur 4.5.445.18 software, and there was a single full-scan mass spectrum in the orbitrap (350-1600 m/z, 60,000 resolution) followed by 2 seconds data-dependent MS/MS scans in an Ion Routing Multipole at 30 normalized collision energy (HCD).

The MS data were analyzed using the MaxQuant algorithm (v.2.4.9.0). Spectra were searched against the *E. coli* K12 protein sequences in the Uniprot database. The global false discovery rate (FDR) cutoff for peptide and protein identification was set to 0.01. The label-free quantification (LFQ) algorithm in MaxQuant was used to compare samples.

Proteomics statistical analysis was performed using the Perseus algorithm (v.2.0.11.0). LFQ intensities for the protein group table were uploaded to Perseus. Protein groups were filtered to retain only proteins with at least 2 valid values in at least one group. Missing values were replaced from a normal distribution (width = 0.3, down shift =1.8). To compare protein intensities, a *t*-test was performed with the correction for multiple hypotheses by FDR-adjusted *p* < 0.05.

### Protein purification

All proteins were recombinantly expressed in *E. coli* BL21(DE3) cells and purified using a Nickel NTA column. For purification of PheS, EcPheS-YFP, PheT, PheT^D1^, DnaA-mCherry, mCherry, YFP, KpPheS-YFP, and BsPheS-YFP, a C-terminal His_6_ tag was introduced. For the purification of mCherry-EcDnaA, mCherry-KpDnaA, mCherry-BsDnaA, and mOrange-HslU, an N-terminal His_6_ tag was introduced. Cultures were grown to an OD_600_ of 0.5 at 37°C, and protein expression was induced overnight at 16°C with 1 mM IPTG. Cells were harvested by centrifugation, resuspended in lysis buffer containing 20 mM Tris-HCl (pH 7.4) and 500 mM NaCl, lysed by sonication, and clarified by centrifugation at 4,000g for 10 min at 4°C. The cleared lysate was applied to a Nickel NTA column, and the resin was washed with buffer containing 20 mM Tris-HCl (pH 7.4), 500 mM NaCl, and 10 mM imidazole. Bound proteins were eluted with the same buffer supplemented with 500 mM imidazole. Eluted fractions were analyzed by SDS-PAGE followed by Coomassie Brilliant Blue staining. Imidazole was removed by repeated buffer exchange using centrifugal ultrafiltration, and the proteins were simultaneously concentrated to ~100 µM into a final buffer containing 20 mM Tris-HCl (pH 7.4) and 500 mM NaCl. Purified proteins were used immediately in subsequent assays, without freezing.

### *In vitro* aggregation assay

Purified proteins in 20 mM Tris-HCl (pH 7.4) and 500 mM NaCl were diluted into assay buffer containing 20 mM Tris-HCl (pH 7.4) and 40 μM thioflavin T (ThT) to the indicated final protein and NaCl concentrations. Samples were loaded into a 96-well plate and sealed to prevent evaporation. The fluorescence intensity of ThT was monitored using a SPARK plate reader with excitation at 435 nm, emission at 490 nm, and 100% gain at room temperature without shaking. Measurements were taken every 10[minutes over 48 hours.

### Statistical analysis

Statistical analyses were performed in GraphPad Prism using tests described in the Figure Legends.

## DATA AVAILABILITY

### Lead contact

Further information and requests for resources and reagents should be directed to, and will be fulfilled by, the lead contact, Jia-Feng Liu.

### Materials availability

This study did not generate new unique reagents. The strains generated and isolated in this study are available upon request.

### Data and code availability

No original code was used in this study.

## ACKNOWLEDGMENTS

We thank Jingjing Wang at the Cell Biology Facility, Center of Biomedical Analysis, Tsinghua University, for help with confocal microscopy. We thank Meng Han in Protein Chemistry and proteomics Facility at Technology Center for Protein Sciences, Tsinghua University, for protein MS analysis. We thank Nathalie Balaban’s lab at Racah Institute of Physics, The Hebrew University of Jerusalem, for providing the *E. coli* KLY strain. This work was supported by the National Key Research and Development Program of China (2022YFC2303202), the Tsinghua-Peking Joint Center for Life Sciences (20111770319), and the Tsinghua University Dushi Program (20251080040) to J. L.

## AUTHOR CONTRIBUTIONS

Conceptualization: W.F.Z., L.J.F.; Methodology: W.F.Z., Z.Y.W., L.J.F.; Investigation: W.F.Z., Z.Y.W., L.J.F.; Visualization: W.F.Z., Z.Y.W.; Funding acquisition: L.J.F.; Project administration: L.J.F.; Supervision: L.J.F.; Writing – original draft: W.F.Z; Writing – review & editing: W.F.Z., Z.Y.W., L.J.F.

## SUPPLEMENTARY INFORMATION

The Supplementary Information includes Extended Data Fig. 1–12 and Supplementary Table 1–5.

