## Extended Data Fig. 1-12 and Supplementary Table 1-5 for "A proteostasis clock tunes bacterial dormancy by timing replication initiation"

### Supplementary INFORMATION

The Supplementary Information includes Extended Data Fig. 1–12 and Supplementary Table 1–5.

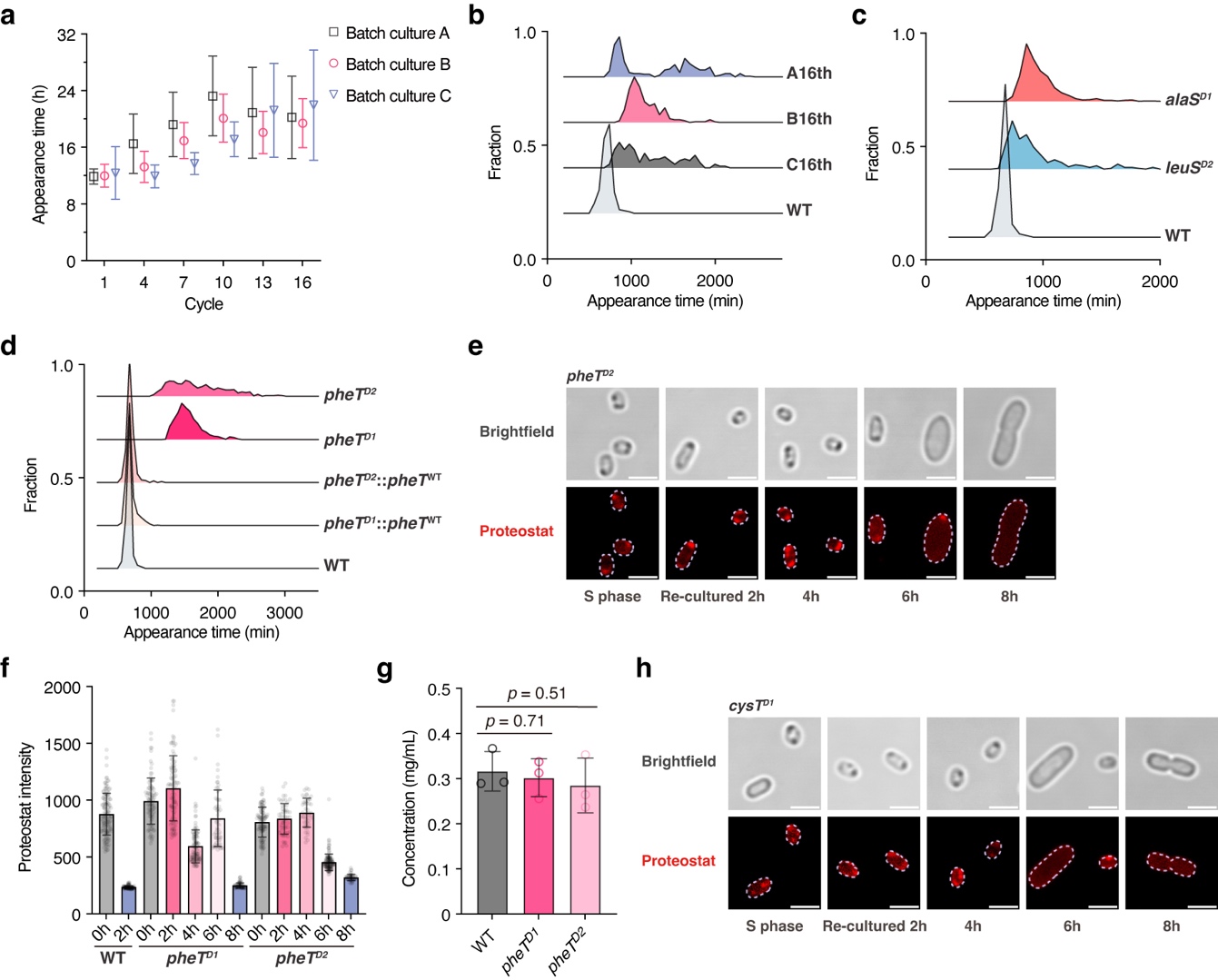

### Extended DATA figures

#### Extended Data Fig. 1. Delayed aggregate disassembly in different dormant mutants.

**a**, Appearance time across cycles in three parallel batch cultures measured by ScanLag. The exposure time for ertapenem (1 μg/mL) was set to 4h and 12h alternately. Data are presented as mean ± SD. Sample sizes N = 234, 209, 189, 222, 161, and 123 for batch culture A. Sample sizes N = 202, 364, 211, 235, 312, and 230 for batch culture B. Sample sizes N = 93, 498, 417, 329, 187, and 253 for batch culture C.

**b**, Appearance time distribution for three evolved batch cultures at the 16th cycle. Sample sizes N = 989, 123, 230, and 253, respectively.

**c**, Appearance time distribution for WT, *leuS^D1^*, and *alaS^D1^* cells measured by ScanLag. Bacteria were sampled from 48-h cultures. Sample sizes N = 989, 262, and 766, respectively.

**d**, Appearance time distribution for WT, *pheT* mutants, and *pheT* mutants with restored *pheT^WT^*. Sample sizes N = 989, 797, 684, 810, and 678, respectively.

**e**, Representative brightfield and fluorescence images of *pheT^D2^* cells stained with Proteostat. Cells from 48-hour cultures were re-cultured in fresh LB medium and sampled at the indicated time points. Scale bars, 2 μm.

**f**, Single-cell Proteostat intensity of WT, *pheT^D1^*, and *pheT^D2^* cells during recovery in fresh LB medium. Data are presented as mean ± SD. Sample sizes N = 130, 95, 92, 98, 108, 59, 64, 100, 54, 45, 141, and 49, respectively.

**g**, Quantification of insoluble protein concentrations in WT and *pheT* mutants using the BCA assay. Data are presented as mean ± SD (n = 3 biological replicates). *P* values were calculated using unpaired two-tailed t-tests.

**h**, Representative brightfield and fluorescence images of *cysT^D1^* cells stained with Proteostat. Cells from 48-hour cultures were re-cultured in fresh LB medium and sampled at the indicated time points. Scale bars, 2 μm.

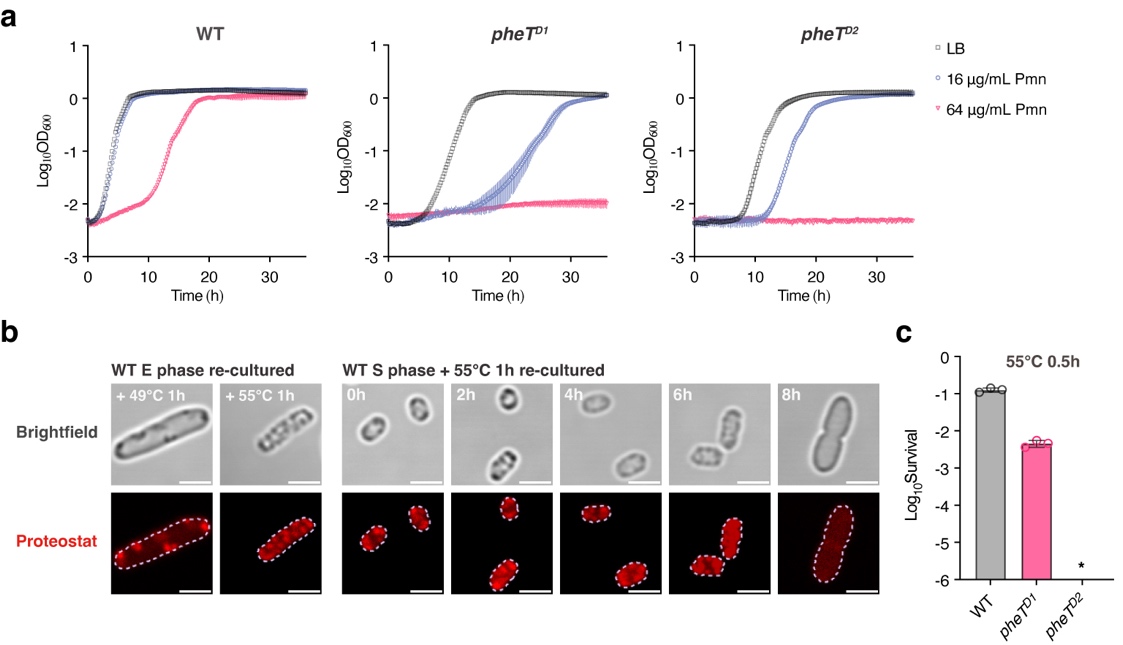

#### Extended Data Fig. 2. Effects of puromycin and heat shock on the growth of WT and pheT mutants.

**a**, Growth curves of WT, *pheT^D1^*, and *pheT^D2^* cells in LB medium supplemented with the indicated concentration of puromycin. Data are presented as mean ± SD (n = 3 biological replicates).

**b**, Representative brightfield and fluorescence images of stress-treated WT cells stained with Proteostat. Left, exponential-phase WT cells were treated at 49 °C or 55 °C for 1h in LB medium. Right, stationary-phase WT cells were treated at 55 °C for 1h and then transferred to fresh LB at 37 °C for regrowth. Scale bars, 2 μm.

**c**, Survival of stationary-phase WT, *pheT^D1^*, and *pheT^D2^* cells after heat shock at 55 °C for 0.5h. Asterisk indicates values below the detection limit (< 10^-7^). Data are presented as mean ± SD (n = 3 biological replicates).

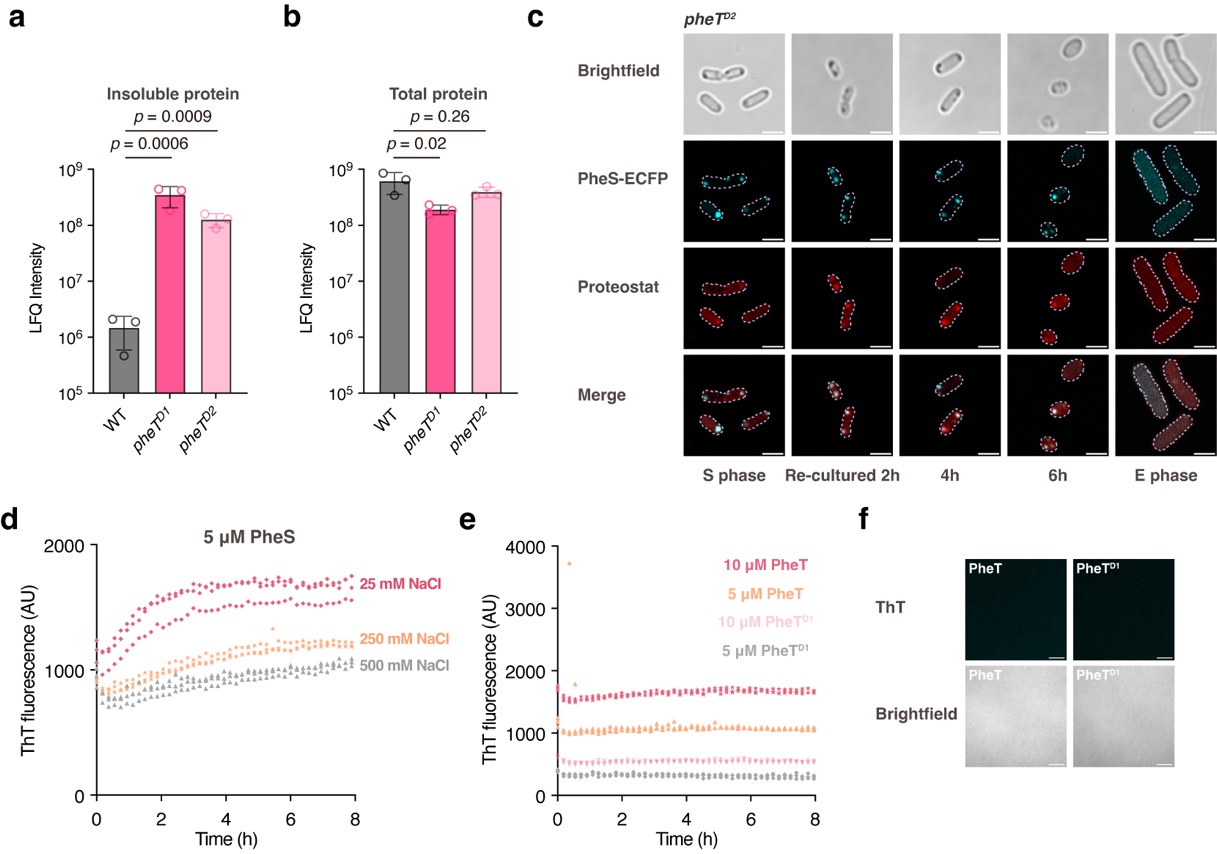

#### Extended Data Fig. 3. Increased PheS aggregation in the pheT mutants.

**a**, **b**, LFQ intensity of PheS in the insoluble proteome (**a**) and total proteome (**b**). Statistical significance was calculated using Student’s *t*-test with the correction to multiple hypotheses by FDR adjusted *p* < 0.05. Data are presented as mean ± SD (n = 3 biological replicates).

**c**, Representative brightfield and fluorescence images of *pheT^D2^* cells expressing PheS-ECFP and stained with Proteostat. Cells from the stationary phase were re-cultured in fresh LB medium and stained with Proteostat at the indicated time points. Scale bars, 2 μm.

**d**, Fluorescence ThT signal over time for 5 μM PheS. Purified PheS was incubated in buffer containing 20 mM Tris at pH 7.4 and the indicated concentration of NaCl. Each point represents the normalized ThT fluorescence of one independent sample at each time point. n = 3 independent samples.

**e**, Fluorescence ThT signal over time for PheT and PheT^D1^. Purified proteins were incubated in buffer containing 20 mM Tris at pH 7.4 and the corresponding NaCl concentrations of 38 mM, 19 mM, 21 mM, and 10.5 mM for 10 μM PheT, 5 μM PheT, 10 μM PheT^D1^, and 5 μM PheT^D1^, respectively. Each point represents the normalized ThT fluorescence of one independent sample at each time point. n = 3 independent samples.

**f**, Representative brightfield and ThT fluorescence images of 5 μM PheT and PheT^D1^ after 48h incubation. Scale bars, 20 μm.

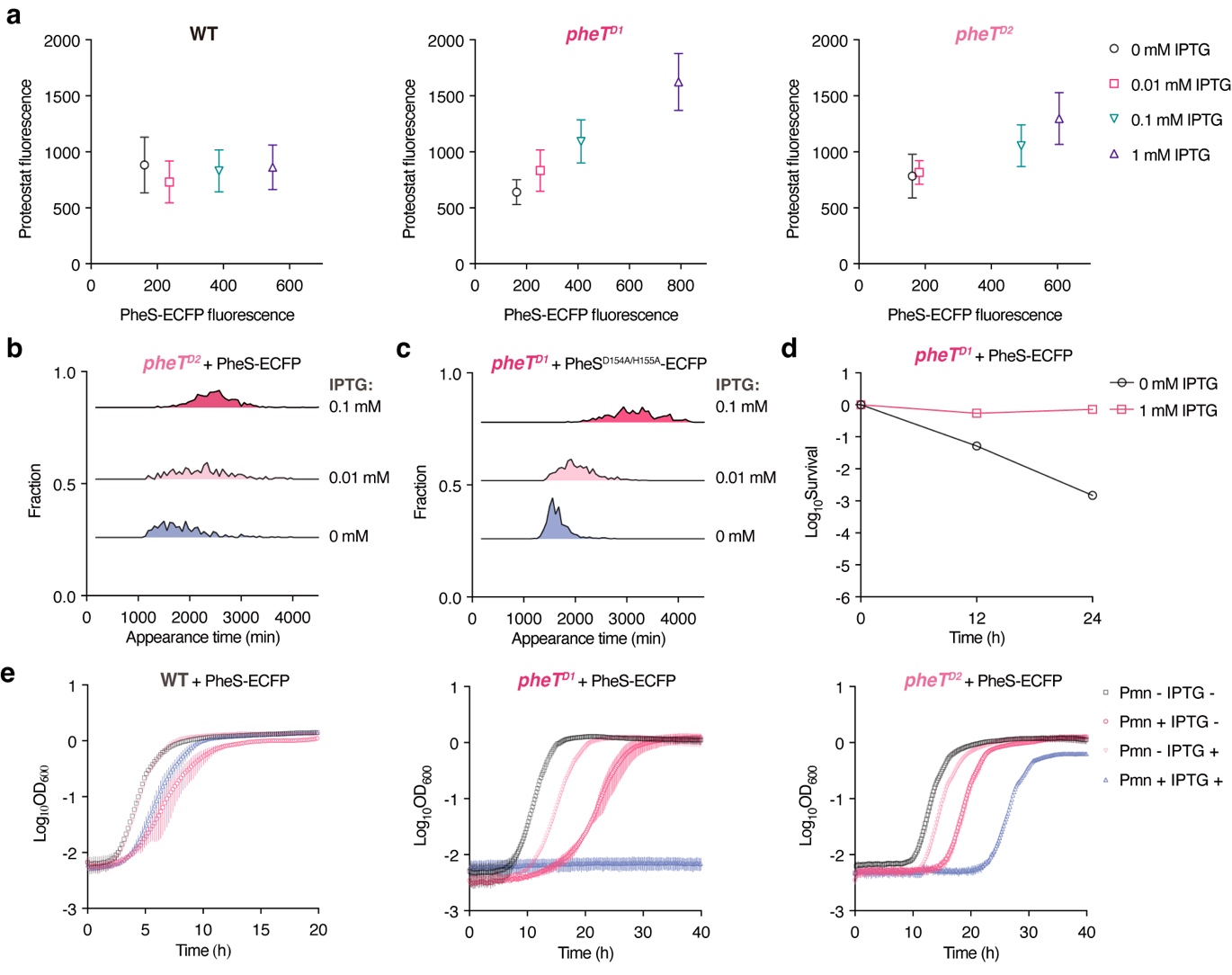

#### Extended Data Fig. 4. Increased PheS aggregation disrupts proteostasis and extends lag time in the pheT mutants.

**a**, Mean single-cell Proteostat fluorescence intensity plotted against the mean single-cell PheS-ECFP intensity of WT, *pheT^D1^*, and *pheT^D2^* cells. Cells carrying an IPTG-inducible PheS-ECFP construct were grown for 48h at the indicated IPTG concentrations. PheS-ECFP and Proteostat fluorescence were quantified separately under the indicated conditions. Data are presented as mean ± SD. Sample sizes N = 176, 220, 197, 167, 459, 184, 169, 131, 110, 82, 312, and 351 for ECFP fluorescence. Sample sizes N = 238, 178, 212, 108, 55, 79, 75, 68, 254, 110, 325, and 161 for Proteostat fluorescence.

**b**, Appearance time distribution for *pheT^D2^* cells after induction of PheS-ECFP expression for 48h at the indicated IPTG concentrations. Sample sizes N = 327, 175, and 704, respectively.

**c**, Appearance time distribution for *pheT^D1^* cells after induction of PheS^D154A/H155A^-ECFP expression for 48h at the indicated IPTG concentrations. Sample sizes N = 691, 668, and 523, respectively.

**d**, Survival of *pheT^D1^* cells after 12 h or 24 h of treatment with 1 μg/mL ertapenem, with or without prior induction of PheS-ECFP expression. Data are presented as mean ± SD (n = 3 biological replicates).

**e**, Growth curves of WT, *pheT^D1^*, and *pheT^D2^* cells under the indicated conditions, including induction of PheS-ECFP expression or puromycin supplementation. IPTG was used at 1 mM for all experiments. Puromycin was used at 32 μg/mL for WT and 16 μg/mL for the *pheT* mutants. Data are presented as mean ± SD (n = 3 biological replicates).

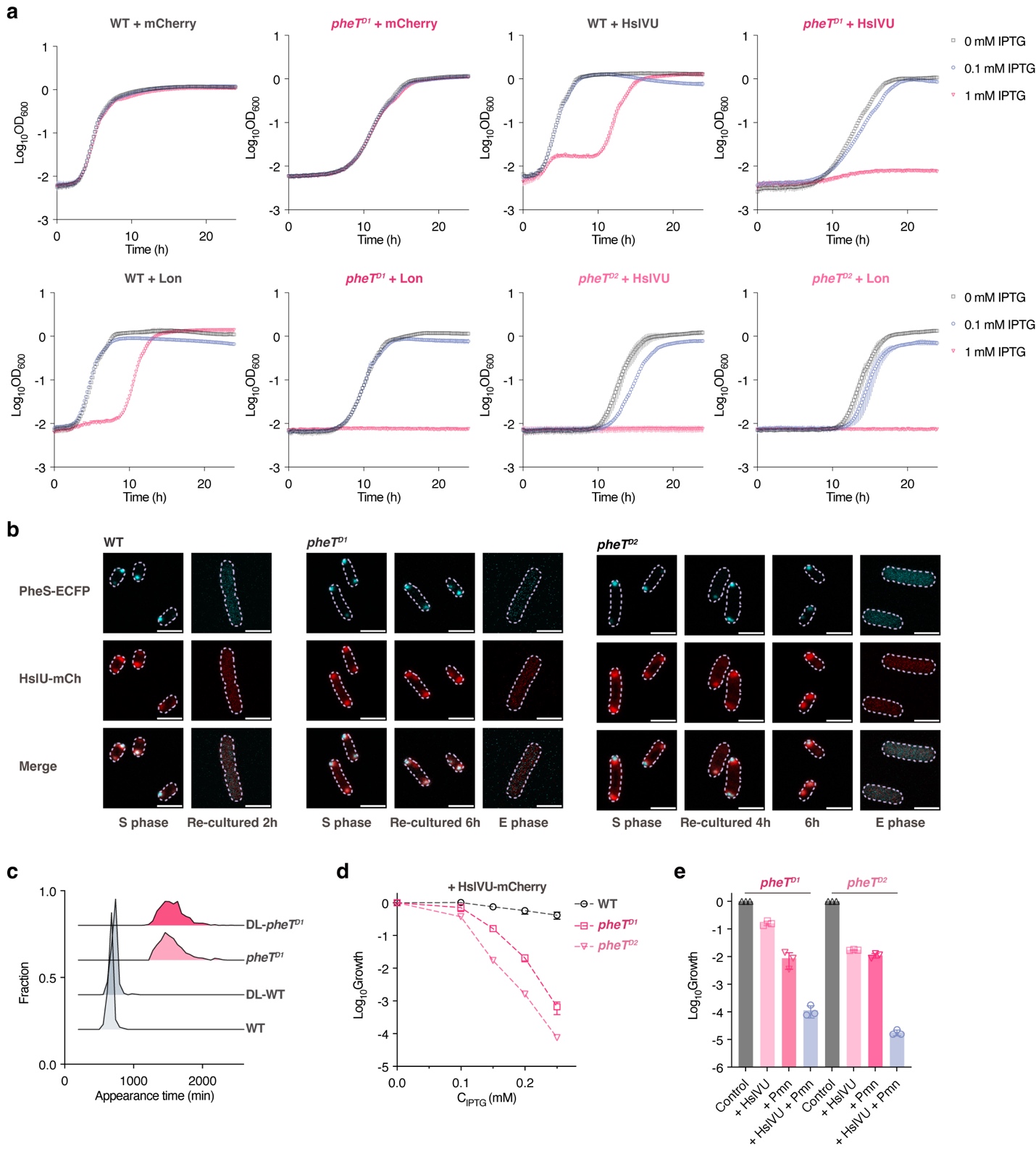

#### Extended Data Fig. 5. Effects of chaperones and proteases on bacterial dormancy exit.

**a**, Growth curves of WT, *pheT^D1^*, and *pheT^D2^* cells during induced expression of proteases with the indicated concentrations of IPTG. Data are presented as mean ± SD (n = 3 biological replicates).

**b**, Representative fluorescence images of WT, *pheT^D1^*, and *pheT^D2^* cells simultaneously expressing HslU-mCherry and PheS-ECFP. Cells from 48-hour cultures were re-cultured in fresh LB medium and sampled at the indicated time points. Scale bars, 2 μm.

**c**, Appearance time distribution for WT, *pheT^D1^*, and dual-labelled (DL) strains (chromosomally labelled HslU-ECFP and mCherry-DnaA). Sample sizes N = 989, 768, 810, and 450, respectively.

**d**, Relative colony-forming fraction of WT, *pheT^D1^*, and *pheT^D2^* cells during induced expression of HslVU with 0.25 mM IPTG, normalized to IPTG-free plates. Data are presented as mean ± SD (n = 3 biological replicates).

**e**, Relative colony-forming fraction of *pheT^D1^* and *pheT^D2^* cells during induced expression of HslVU with 0.15 mM IPTG or 16 μg/mL puromycin supplementation, normalized to drug-free plates. Data are presented as mean ± SD (n = 3 biological replicates).

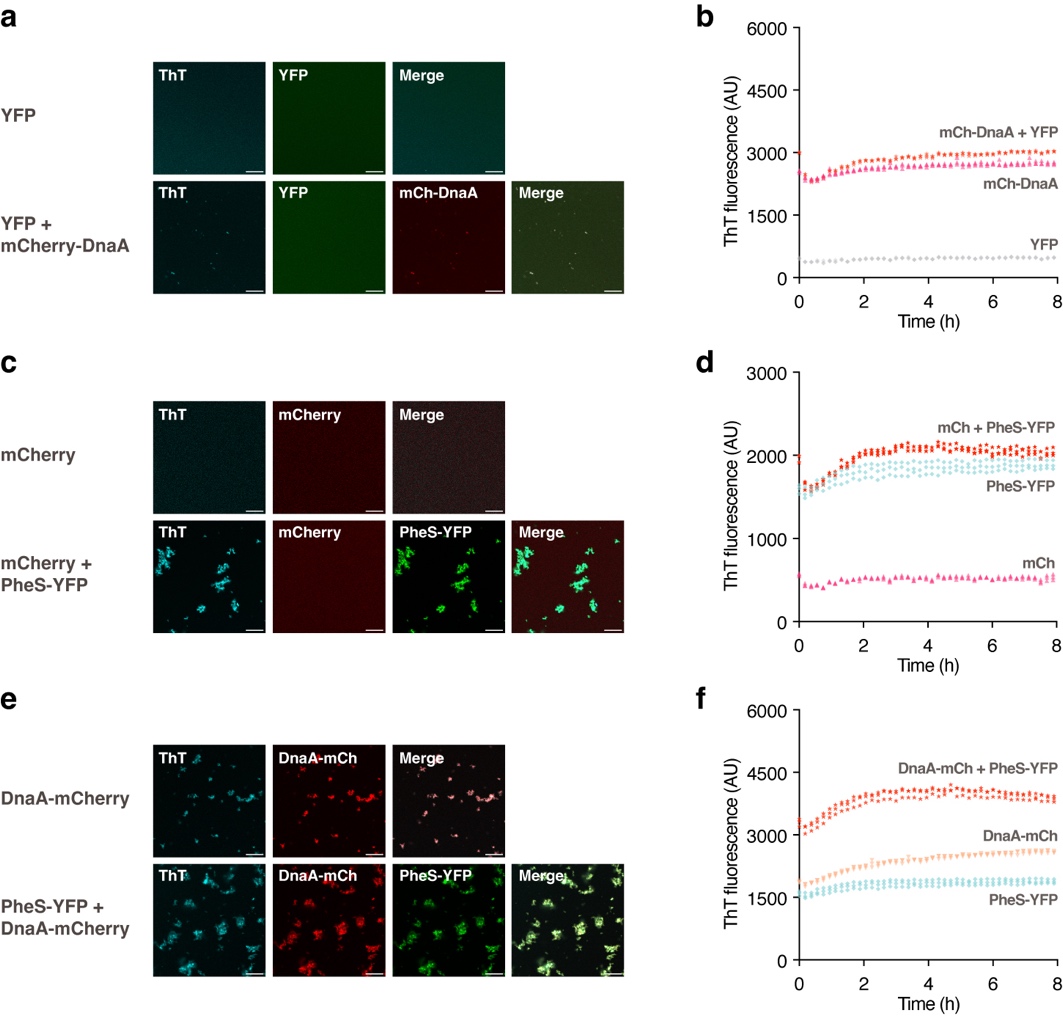

#### Extended Data Fig. 6. In vitro reconstitution of protein aggregates.

**a**, **c**, **e**, Representative fluorescence images of the indicated samples after 48h incubation. Each protein was used at 5 μM in buffer containing 20 mM Tris at pH 7.4 and 62 mM NaCl. Scale bars, 20 μm.

**b**, **d**, **f**, Fluorescence ThT signal over time during incubation of the indicated protein samples. Each protein was used at 5 μM in buffer containing 20 mM Tris at pH 7.4 and 62 mM NaCl. Each point represents the normalized ThT fluorescence of one independent sample at each time point. n = 3 independent samples.

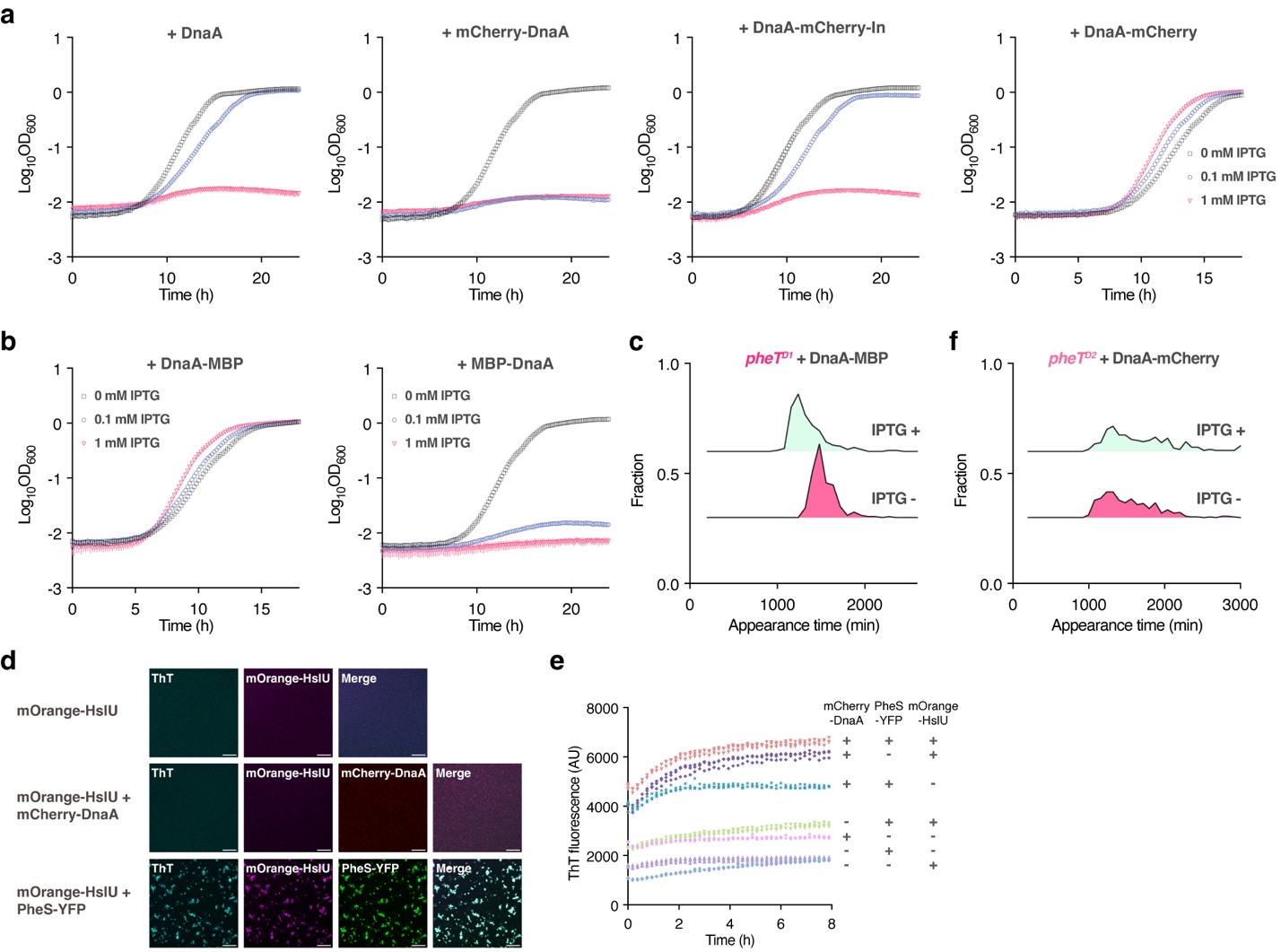

#### Extended Data Fig. 7. Soluble DnaA variant facilitates bacterial exit from dormancy.

**a**, Growth curves of *pheT^D1^* cells expressing mCherry-tagged DnaA variants with the indicated concentrations of IPTG. Data are presented as mean ± SD (n = 3 biological replicates).

**b**, Growth curves of *pheT^D1^* cells expressing MBP-tagged DnaA variants with the indicated concentrations of IPTG. DnaA-MBP, DnaA with carboxy-terminus labelled MBP; MBP-DnaA, DnaA with amino-terminus labelled MBP. Data are presented as mean ± SD (n = 3 biological replicates).

**c**, Appearance time distribution for *pheT^D1^* cells during induced expression of DnaA-MBP with 0.25 mM IPTG. Sample sizes N = 203 and 211, respectively.

**d**, Representative fluorescence images of the indicated samples after 48h incubation. Each protein was used at 5 μM in buffer containing 20 mM Tris at pH 7.4 and 62 mM NaCl. Scale bars, 20 μm.

**e**, Fluorescence ThT signal over time during incubation of the indicated protein samples. Each protein was used at 5 μM in buffer containing 20 mM Tris at pH 7.4 and 62 mM NaCl. Each point represents the normalized ThT fluorescence of one independent sample at each time point. n = 3 independent samples.

**f**, Appearance time distribution for *pheT^D2^* cells during induced expression of DnaA-mCherry with 0.25 mM IPTG. Sample sizes N = 343 and 200, respectively.

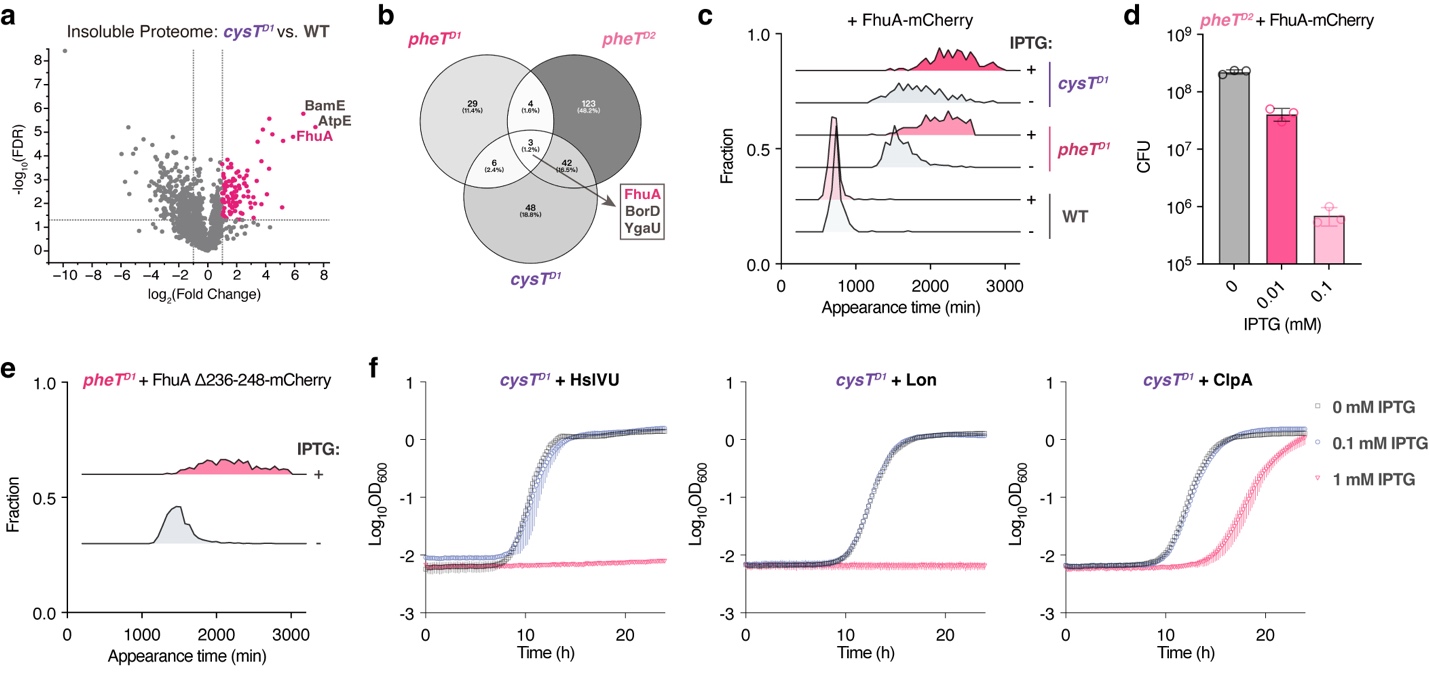

#### Extended Data Fig. 8. Disrupted proteostasis in the long-lag cysT^D1^ mutant.

**a**, Volcano plots showing MS analysis of the insoluble proteome in *cysT^D1^* relative to WT at the stationary phase. Student’s *t*-test with the correction to multiple hypotheses by FDR adjusted *p* < 0.05. n = 3 biological replicates.

**b**, Venn diagram showing overlap among proteins with increased aggregation in *pheT^D1^*, *pheT^D2^*, and *cysT^D1^* relative to WT.

**c**, Appearance time distribution for WT, *pheT^D1^*, and *cysT^D1^* cells after induction of FhuA-mCherry expression for 48h with 0.1 mM IPTG. Sample sizes N = 228, 332, 383, 219, 244, and 110, respectively.

**d**, CFU of *pheT^D2^* cells after induction of FhuA-mCherry expression for 48 h at the indicated IPTG concentrations. Data are presented as mean ± SD (n = 3 biological replicates).

**e**, Appearance time distribution for *pheT^D1^* cells after induction of FhuA Δ236-248-mCherry expression for 48h with 0.1 mM IPTG. Sample sizes N = 1185 and 368, respectively.

**f**, Growth curves of *cysT^D1^* cells during induced expression of proteases with the indicated concentrations of IPTG. Data are presented as mean ± SD (n = 3 biological replicates).

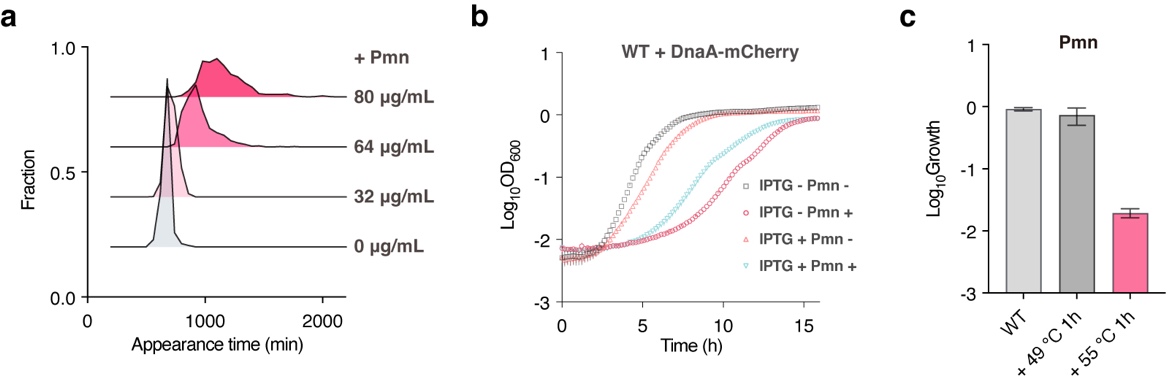

#### Extended Data Fig. 9. Effect of proteotoxic stress on bacterial lag time.

**a**, Appearance time distribution for WT cells incubated with the indicated concentrations of puromycin. Sample sizes N = 989, 961, 805, and 557, respectively.

**b**, Growth curves of WT cells under the indicated conditions, including inducible DnaA-mCherry expression or supplementation with 64 μg/mL puromycin. Data are presented as mean ± SD (n = 3 biological replicates).

**c**, Relative colony-forming fraction of stress-treated WT cells on puromycin plates (64 μg/mL), normalized to the corresponding IPTG-free controls. Data are presented as mean ± SD (n = 3 biological replicates).

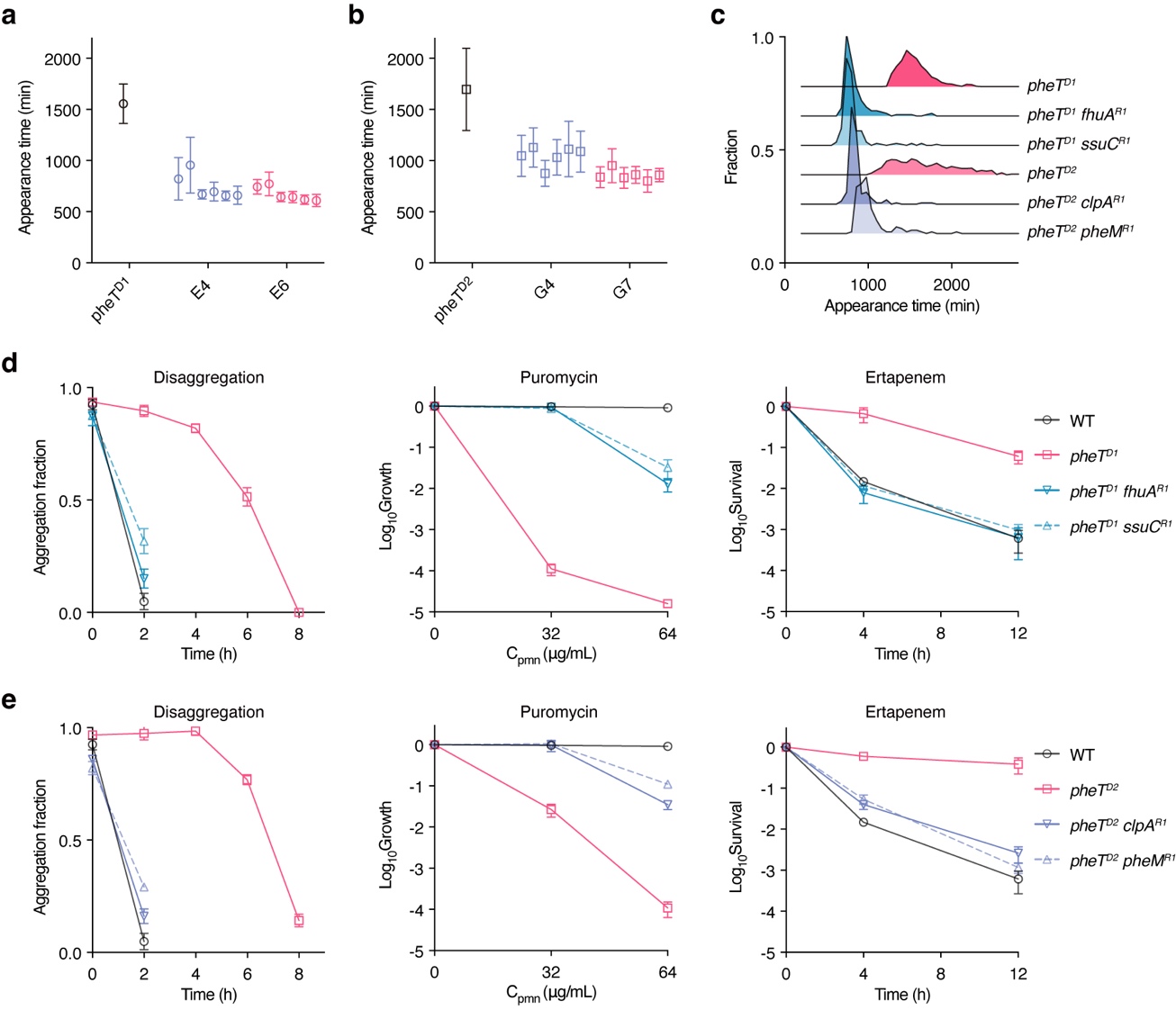

#### Extended Data Fig. 10. Restored proteostasis and shortened lag time in reverse mutants.

**a**, **b**, Appearance time across cycles in six parallel batch cultures during reverse evolution of *pheT^D1^* (**a**) and *pheT^D2^* (**b**), measured by ScanLag. Dormant mutants were serially passaged in antibiotic-free LB medium. E4, the fourth cycle; E6, the sixth cycle; E7, the seventh cycle. Data are presented as mean ± SD. Sample sizes N = 810, 271, 282, 369, 434, 461, 395, 451, 321, 357, 427, 431, and 390 in (**a**). Sample sizes N = 678, 193, 221, 345, 278, 125, 114, 312, 206, 149, 189, 162, and 253 in (**b**).

**c**, Appearance time distribution for the indicated reverse mutants. Sample sizes N = 136, 151, 678, 202, 173, and 810, respectively.

**d**, **e**, The disaggregation dynamics and antibiotic sensitivity of reverse mutants. Left, quantification of the fraction of cells containing Proteostat-positive aggregates upon re-culturing. Data are presented as mean ± SD (n = 4 biological replicates). Middle, relative colony-forming fraction of reverse mutants on solid medium containing the indicated concentrations of puromycin, normalized to drug-free plates. Data are presented as mean ± SD (n = 3 biological replicates). Right, survival of reverse mutants after 4h or 12h of 1 µg/mL ertapenem treatment. Data are presented as mean ± SD (n = 3 biological replicates).

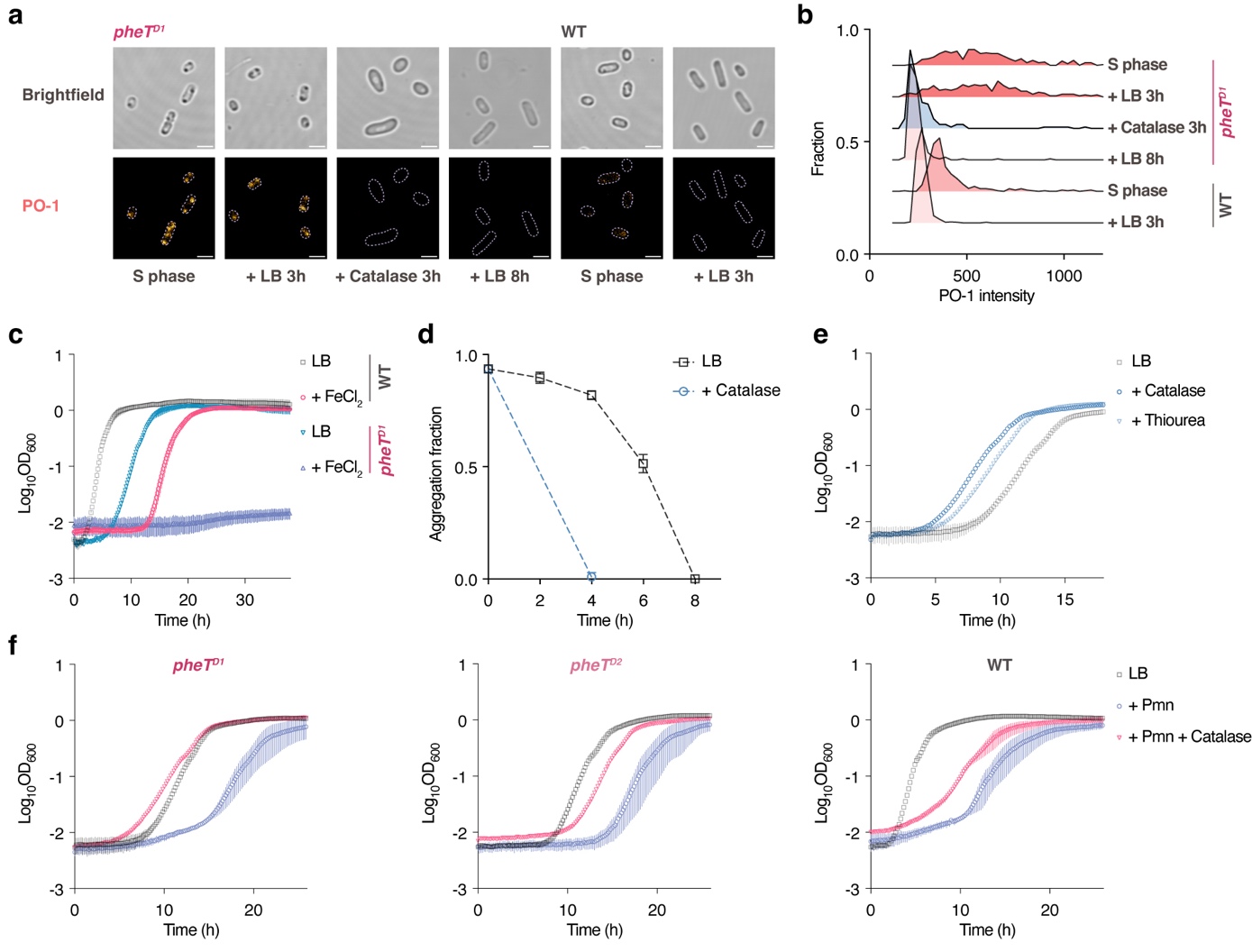

#### Extended Data Fig. 11. Hydroxyl radical scavenging facilitates the recovery of proteostasis and the exit from dormancy.

**a**, Representative brightfield and fluorescence images of WT and *pheT^D1^* cells stained with Peroxy Orange 1 (PO-1). Cells from the stationary phase were re-cultured in LB or LB supplemented with 0.1 mg/mL catalase and sampled at the indicated time points. Scale bars, 2 μm.

**b**, PO-1 fluorescence intensity distribution for WT and *pheT^D1^* cells at the indicated time points during recovery. Sample sizes N = 445, 374, 277, 166, 243, and 269, respectively.

**c**, Growth curves of WT and *pheT^D1^* cells in LB supplemented with 6.8 mM FeCl_2_. Data are presented as mean ± SD (n = 3 biological replicates).

**d**, Quantification of the fraction of *pheT^D1^* cells containing Proteostat-positive aggregates upon re-culturing in LB or LB supplemented with 0.1 mg/mL catalase. Data are presented as mean ± SD (n = 4 biological replicates).

**e**, Growth curves of *pheT^D1^* cells in LB supplemented with 0.1 mg/mL catalase or 30 mM thiourea. Data are presented as mean ± SD (n = 3 biological replicates).

**f**, Growth curves of WT, *pheT^D1^*, and *pheT^D2^* cells under the indicated conditions, including puromycin and catalase supplementation. Puromycin was used at 64 μg/mL for WT and 16 μg/mL for *pheT^D1^* and *pheT^D2^* cells. Catalase was used at 0.4 mg/mL for WT, 0.1 mg/mL for *pheT^D1^*, and 0.2 mg/mL for *pheT^D2^*. Data are presented as mean ± SD (n = 3 biological replicates).

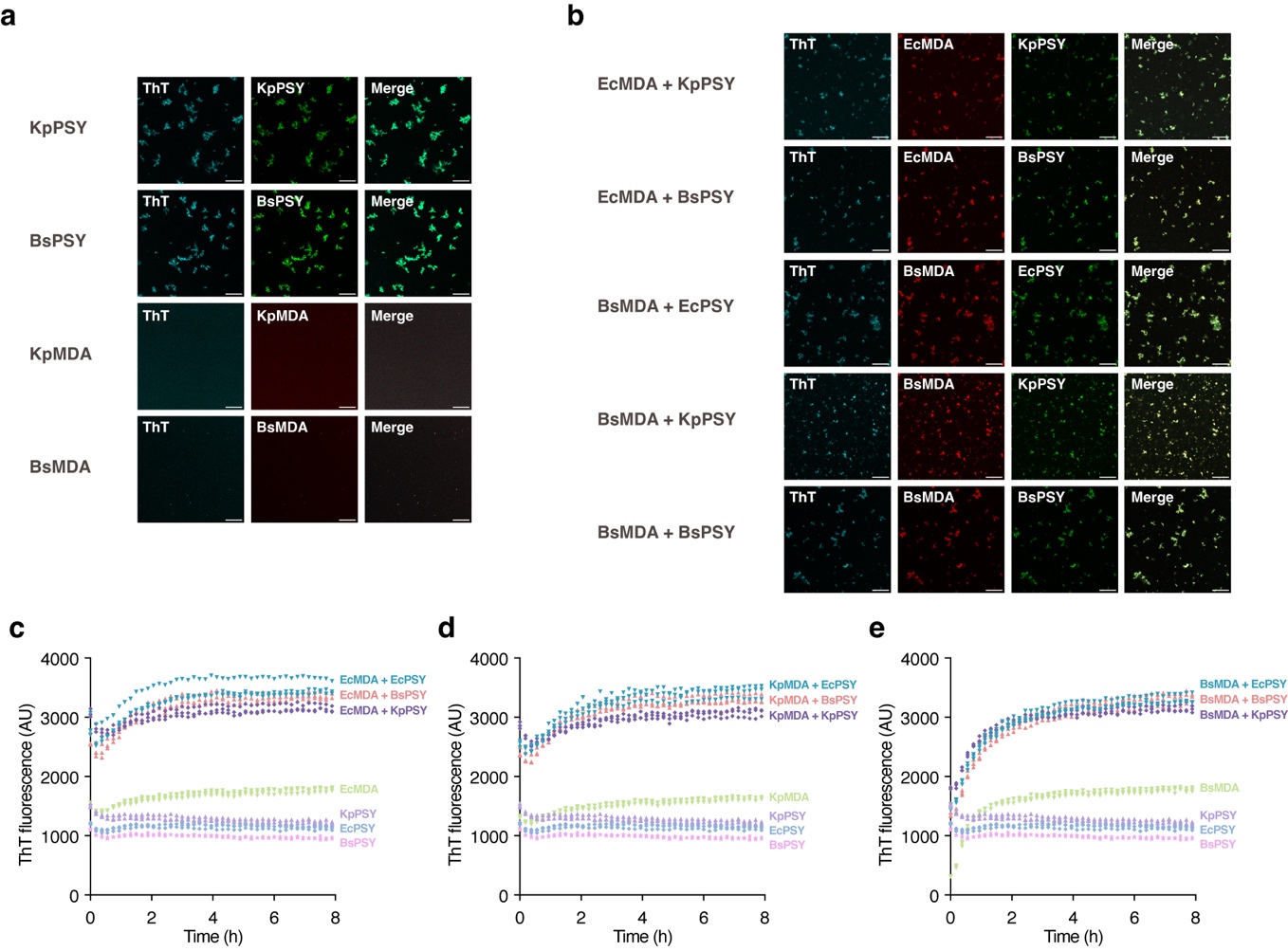

#### Extended Data Fig. 12. Co-aggregation of PheS and DnaA from different bacterial species.

**a**, Representative fluorescence images of purified PheS-YFP (PSY) and mCherry-DnaA (MDA) from *K. pneumoniae* and *B. subtilis* after 48h incubation alone. Each protein was used at 5 μM in buffer containing 20 mM Tris at pH 7.4 and 62 mM NaCl. Scale bars, 20 μm.

**b**, Representative brightfield and fluorescence images of PSY and MDA from the indicated species after 48h co-incubation. Each protein was used at 5 μM in buffer containing 20 mM Tris at pH 7.4 and 62 mM NaCl. Scale bars, 20 μm.

**c**–**e**, Fluorescence ThT signal over time during incubation of the indicated protein samples. Each protein was used at 5 μM in buffer containing 20 mM Tris at pH 7.4 and 62 mM NaCl. Each point represents the normalized ThT fluorescence of one independent sample at each time point. n = 3 independent samples.

### Supplemental tables

#### Supplementary Table 1. Mutations identified by whole genome sequencing and verified by Sanger sequencing under the evolution of ertapenem

| Strain | Genomic position | Mutation | Amino Acid Substitution | Gene | Annotation |
| --- | --- | --- | --- | --- | --- |
| *alaS^D1^* | 2795904 | T>G | 26 AA insertion | *alaS* | Alanine-tRNA ligase |
| *leuS^D1^* | 671398 | A>G | V806A | *leuS* | Leucine-tRNA ligase |
| *cysT^D1^* | 1967629 | T>G | none | *cysT* | Transfer RNA of cysteine |
| *pheT^D1^* | 1771783 | Δ1bp | truncation | *pheT* | Phenylalanine-tRNA ligase subunit β |
| *pheT^D2^* | 1772615-1772668 | Δ54bp | 18 AA deletion | *pheT* | Phenylalanine-tRNA ligase subunit β |
|  | 2812864 | A>C | S362A | *hypF* | Carbamoyltransferase |
|  | 487888 | A>G | none | *mscK* | Mechanosensitive channel |

#### Supplementary Table 2. Additional reverse mutations in the pheT mutants identified by whole genome sequencing and verified by Sanger sequencing

| Strain | Genomic position | Mutation | Amino Acid Substitution | Gene | Annotation |
| --- | --- | --- | --- | --- | --- |
| *pheT^D1^ fhuA^R1^* | 167762 | Δ1bp | truncation | *fhuA* | Ferrichrome outer membrane transporter/phage receptor |
| *pheT^D1^ ssuC^R1^* | 993820 | C>T | truncation | *ssuC* | Aliphatic sulfonate ABC transporter membrane subunit |
| *pheT^D2^ clpA^R1^* | 923318 | C>T | truncation | *clpA* | ATP-dependent protease specificity component and chaperone |
|  | 2280252 | T>C | none | *-* | Non-coding region |
| *pheT^D2^ pheM^R1^* | 1774449-1774502 | Δ54bp | none | *pheM* | Phenylalanine-tRNA ligase operon leader peptide *pheM* terminator |

#### Supplementary Table 3. Strains

| Strain Name | Description | Reference |
| --- | --- | --- |
| ***E. coli*** |  |  |
| KLY | KL16-YFP Cam | Balaban *et al*. |
| KLY *metG^T^* | KLY evolution | Levin-Reisman *et al*. |
| *pheT^D1^* | KLY evolution (see Table S1) | This work |
| *pheT^D2^* | KLY evolution (see Table S1) | This work |
| *alaS^D1^* | KLY evolution (see Table S1) | This work |
| *leuS^D1^* | KLY evolution (see Table S1) | This work |
| *cysT^D1^* | KLY evolution (see Table S1) | This work |
| *pheT^D1^ fhuA^R1^* | *pheT^D1^* reverse evolution (see Table S2) | This work |
| *pheT^D1^ ssuC^R1^* | *pheT^D1^* reverse evolution (see Table S2) | This work |
| *pheT^D2^ clpA^R1^* | *pheT^D2^* reverse evolution (see Table S2) | This work |
| *pheT^D2^ pheM^R1^* | *pheT^D2^* reverse evolution (see Table S2) | This work |
| *pheT^D1^*::*pheT^WT^* | Restoration of the WT *pheT* in *pheT^D1^* through P1 transduction | This work |
| *pheT^D2^*::*pheT^WT^* | Restoration of the WT *pheT* in *pheT^D2^* through P1 transduction | This work |
| WT::*hslVU-ECFP & mCherry-dnaA* | Label HslVU and DnaA with fluorescent proteins in WT with λ Red recombineering, mCherry at the N-terminus | This work |
| *pheT^D1^*::*hslVU-ECFP & mCherry-dnaA* | Label HslVU and DnaA with fluorescent proteins in *pheT^D1^* with λ Red recombineering, mCherry at the N-terminus | This work |
| *pheT^D1^*::*hslVU-ECFP & dnaA-mCherry* | Label HslVU and DnaA with fluorescent proteins in *pheT^D1^* with λ Red recombineering, mCherry at the C-terminus | This work |
| *pheT^D1^*::*hslVU-ECFP & dnaA-mCherry-In* | Label HslVU and DnaA with fluorescent proteins in *pheT^D1^* with λ Red recombineering, mCherry is at the internal sites of DnaA | This work |
| ***K. pneumoniae*** |  |  |
| ATCC 43816 |  |  |
| S837 | Isolated from hospital | This work |
| ***B. subtilis*** |  |  |
| PY79 |  |  |

#### Supplementary Table 4. Plasmids

| Name | Description | Reference |
| --- | --- | --- |
| pET28a-pheS-ECFP | For the inducible expression of PheS-ECFP with IPTG | This work |
| pET28a-pheS^D154A/H155A^-ECFP | For the inducible expression of PheS^D154A/H155A^-ECFP with IPTG | This work |
| pET28a-pheS | For the inducible expression of PheS with IPTG | This work |
| pET28a-ECFP | For the inducible expression of ECFP with IPTG | This work |
| pET28a-mCherry | For the inducible expression of mCherry with IPTG | This work |
| pET28a-dnaK-mCherry | For the inducible expression of DnaK-mCherry with IPTG | This work |
| pET28a-clpB-mCherry | For the inducible expression of ClpB-mCherry with IPTG | This work |
| pET28a-tig-mCherry | For the inducible expression of Tig-mCherry with IPTG | This work |
| pET28a-ibpA-mCherry | For the inducible expression of IbpA-mCherry with IPTG | This work |
| pET28a-clpS-mCherry | For the inducible expression of ClpS-mCherry with IPTG | This work |
| pET28a-hslVU-mCherry | For the inducible expression of HslVU-mCherry with IPTG | This work |
| pET28a-lon-mCherry | For the inducible expression of Lon-mCherry with IPTG | This work |
| pET28a-clpA-mCherry | For the inducible expression of ClpA-mCherry with IPTG | This work |
| pET28a-clpX-mCherry | For the inducible expression of ClpX-mCherry with IPTG | This work |
| pET28a-clpP-mCherry | For the inducible expression of ClpP-mCherry with IPTG | This work |
| pET28a-ftsH-mCherry | For the inducible expression of FtsH-mCherry with IPTG | This work |
| pET28a-dnaA | For the inducible expression of DnaA with IPTG | This work |
| pET28a-dnaA-mCherry | For the inducible expression of DnaA-mCherry with IPTG | This work |
| pET28a-mCherry-dnaA | For the inducible expression of mCherry-DnaA with IPTG | This work |
| pET28a-dnaA-mCherry-In | For the inducible expression of DnaA-mCherry-In with IPTG | This work |
| pET28a-dnaA-MBP | For the inducible expression of DnaA-MBP with IPTG | This work |
| pET28a-MBP-dnaA | For the inducible expression of MBP-DnaA with IPTG | This work |
| pET28a-aTc-pheS-ara-hslVU-ECFP-lac-dnaA-mCherry | For the inducible expression of PheS with anhydrotetracycline, HslVU-ECFP with arabinose, and DnaA-mCherry with IPTG | This work |
| pET28a-fhuA-mCherry | For the inducible expression of FhuA-mCherry with IPTG | This work |
| pET28a-fhuA Δ236-248-mCherry | For the inducible expression of FhuA Δ236-248-mCherry with IPTG | This work |
| pSC101-Red-Tet | Helper plasmid for λ Red recombineering with IPTG induction | This work |
| pET28a-pheS-His_6_ | For the purification of PheS | This work |
| pET28a-EcpheS-YFP-His_6_ | For the purification of EcPheS-YFP | This work |
| pET28a-pheT-His_6_ | For the purification of PheT | This work |
| pET28a-pheT^D1^-His_6_ | For the purification of PheT^D1^ | This work |
| pET28a-His_6_-mCherry-EcdnaA | For the purification of mCherry-EcDnaA | This work |
| pET28a-His_6_-mOrange-hslU | For the purification of mOrange-HslU | This work |
| pET28a-dnaA-mCherry-His_6_ | For the purification of DnaA-mCherry | This work |
| pET28a-YFP-His_6_ | For the purification of YFP | This work |
| pET28a-mCherry-His_6_ | For the purification of mCherry | This work |
| pET28a-KppheS-YFP-His_6_ | For the purification of KpPheS-YFP | This work |
| pET28a-BspheS-YFP-His_6_ | For the purification of BsPheS-YFP | This work |
| pET28a-His_6_-mCherry-KpdnaA | For the purification of mCherry-KpDnaA | This work |
| pET28a-His_6_-mCherry-BsdnaA | For the purification of mCherry-BsDnaA | This work |

#### Supplementary Table 5. Primers

| Name | Sequence | usage |
| --- | --- | --- |
| lac-alaS-F | aatataccatgggcagcagcatgagcaagagcaccgctga | *alaS^D1^* sequence validation |
| alasmut-R | atgccgccgccgccgccgccacctgactttatcgttg |  |
| lac-leuS-F | aatataccatgggcagcagcatgcaagagcaataccgccc | *leuS^D1^* sequence validation |
| leuSwt-R | atgccgccgccgccgccgccgccaacgaccagattga |  |
| pheT-13 | tggcgaaagagggcgaaacg | *pheT^D1^* and *pheT^D2^* sequence validation |
| phetwt-R | atgccgccgccgccgccgccatccctcaatgatgcct |  |
| lpheTm-F | aatataccatgggcagcagcatgaaattcagtgaactgtg |  |
| hypF-test-F | gcacaacgtattgcggggtaaaatc |  |
| hypF-test-R | gaacatgcggaacaagaggcg |  |
| mscK-test-F | cattgctgcctatccatttctggtc |  |
| mscK-test-R | cttgtaaaccaaaaccaagacctacgg |  |
| cysT-test-F | ccttgtggatctcaactcgc | *cysT^D1^* sequence validation |
| cysT-test-R | acatttgggttgagtacgcc |  |
| fhuA-test-F | atggcgcgttccaaaactgc | *pheT^D1^* *fhuA^R1^* sequence validation |
| fhuA-test-R | ttagaaacggaaggttgcggttg |  |
| ssuC-F | aatataccatgggcagcagcatggcaacgccagtgaagaa | *pheT^D1^* *ssuC^R1^* sequence validation |
| ssuC-R | atgccgccgccgccgccgcctaccgtggcctccttca |  |
| clpA-test-F | cggtaaaaccgcgattgcggaag | *pheT^D2^* *clpA^R1^* sequence validation |
| clpA-test-R | caatgcccaaagctttcgaaagc |  |
| eco-test-F | cgtatctgggcgatgctggaatg |  |
| eco-test-R | gaagtggcggatattgcgttgtaatg |  |
| pheM-test-F | gtcatctgaagggttaagtgccc | *pheT^D2^* *pheM^R1^* sequence validation |
| pheM-test-R | ctactacggtgcgcgttctcgc |  |
| ECFP-F | aatataccatgggcagcagcatgctgagcaagggcgagga | Cloning of *ECFP* into pET28a-ECFP |
| ECFP-R | gctttgttagcagccggatcttacttgtacagctcgtcca |  |
| pheS-F | aatataccatgggcagcagcatgtcacatctcgcagaact | Cloning of *pheS* into pET28a-pheS, pET28a-pheS-ECFP, and pET28a-pheS^D154A/H155A^-ECFP |
| ki-pheS-4 | gccgccgccgccgccgcctttaaactgtttgaggaaac |  |
| pheS-R | gctttgttagcagccggatcttatttaaactgtttgaggaaacgc |  |
| pheS-TA-F | gcagcagacactttctggtttgacac |  |
| pheS-TA-R | agaaagtgtctgctgcagcgcgcgccgggtggtgac |  |
| lac-mch-F | aatataccatgggcagcagcatggtgagcaagggcgagga | Cloning of *mCherry* into pET28a-mCherry |
| lac-mch-R | gctttgttagcagccggatctcacttgtacagctcgtcca |  |
| lac-dnak-F | aatataccatgggcagcagcatgggtaaaataattggtat | Cloning of *dnaK* into pET28a-dnaK-mCherry |
| lac-dnak-R | tcaccatgccgccgccgccgccgccttttttgtctttgacttctt |  |
| clpB-F | aatataccatgggcagcagcatgaacgatcaaggtgctga | Cloning of *clpB* into pET28a-clpB-mCherry |
| clpB-R | atgccgccgccgccgccgccctggacggcgacaatccggt |  |
| tig-F | aatataccatgggcagcagcatgcaagtttcagttgaaac | Cloning of *tig* into pET28a-tig-mCherry |
| tig-R | atgccgccgccgccgccgcccgcctgctggttcatcagct |  |
| ibpA-F | aatataccatgggcagcagcatgcgtaactttgatttatc | Cloning of *ibpA* into pET28a-ibpA-mCherry |
| ibpA-R | atgccgccgccgccgccgccgttgatttcgatacggcgcg |  |
| clpS-F | aatataccatgggcagcagcatgggtaaaacgaacgactg | Cloning of *clpS* into pET28a-clpS-mCherry |
| clpS-R | atgccgccgccgccgccgccggctttttctagcgtacaca |  |
| hslV-F | aatataccatgggcagcagcgtgacaactatagtaagcgt | Cloning of *hslVU* into pET28a-hslVU-mCherry |
| hslU-R | atgccgccgccgccgccgcctaggataaaacggctcagat |  |
| lon-F | aatataccatgggcagcagcatgaatcctgagcgttctga | Cloning of *lon* into pET28a-lon-mCherry |
| lon-R | atgccgccgccgccgccgccttttgcagtcacaacct |  |
| lac-clpA-F | aatataccatgggcagcagcatgctcaatcaagaactgga | Cloning of *clpA* into pET28a-clpA-mCherry |
| clpawt-R | atgccgccgccgccgccgccatgcgctgcttccgcct |  |
| clpX-F | aatataccatgggcagcagcatgacagataaacgcaaaga | Cloning of *clpX* into pET28a-clpX-mCherry |
| clpX-R | atgccgccgccgccgccgccttcaccagatgcctgttgcg |  |
| clpP-F | aatataccatgggcagcagcatgtcatacagcggcgaacg | Cloning of *clpP* into pET28a-clpP-mCherry |
| clpP-R | atgccgccgccgccgccgccattacgatgggtcagaa |  |
| ftsH-F | aatataccatgggcagcagcatggcgaaaaacctaatact | Cloning of *ftsH* into pET28a-ftsH-mCherry |
| ftsH-R | atgccgccgccgccgccgcccttgtcgcctaactgctctg |  |
| lac-dnaA-F | aatataccatgggcagcagcgtgtcactttcgctttggca | Cloning of *dnaA* into pET28a-dnaA, pET28a-dnaA-mCherry, pET28a-dnaA-mCherry-In, pET28a-dnaA-MBP, and pET28a-mCherry-dnaA |
| lac-dnaA-R | gctttgttagcagccggatcttacgatgacaatgttctga |  |
| dnaA-R | gccgccgccgccgccgcccgatgacaatgttctgatta |  |
| mch-dnaA-F | caagggcggcggcggcggcggctcactttcgctttggcagcagtg |  |
| mch-dnaA-R | gctttgttagcagccggatcttacgatgacaatgttctgatta |  |
| ki-dma-m1-1 | tatcctcctcgcccttgctcaccatcgtttgcgtcaccggtttggtgccg |  |
| ki-dma-m1-2 | atggtgagcaagggcgaggaggata |  |
| ki-dma-m1-3 | cttgtacagctcgtccatgccgccg |  |
| ki-dma-m1-4 | cggcggcatggacgagctgtacaagcagccgcaacgtgctgcgccttcta |  |
| lac-MBP-F | aatataccatgggcagcagcatgaaaatcgaagaaggtaaactggtaa | Cloning of *MBP* into pET28a-MBP-dnaA, and pET28a-dnaA-MBP |
| dnaA-MBP-F | tcgggcggcggcggcggcggcaaaatcgaagaaggtaaactggta |  |
| dnaA-MBP-R | ttcgggctttgttagcagccggatcttaagtctgcgcgtctttcagggcttcat |  |
| MBP-R | gccgccgccgccgccgccagtctgcgcgtctttcagggcttcatcg |  |
| MBP-dnaA-F | gactggcggcggcggcggcggctcactttcgctttggcagcagtgtc |  |
| MBP-dnaA-R | gctttgttagcagccggatcttacgatgacaatgttctgattaaat |  |
| rha-ECFP-F0 | tgttgctcaggtcgcagacgatttgtcctactcaggagag | Construction of pET28a-aTc-pheS-ara-hslVU-ECFP-lac-dnaA-mCherry |
| bb-rha-R | cgtctgcgacctgagcaaca |  |
| rha-ECFP-R0 | taaatcgatgcaggtggcacttttcggggaaatgtggagg |  |
| rha-ECFP-F00 | gtgccacctgcatcgatttattacttgtacagctcgtcca |  |
| ara-hslV-R0 | ctctaaggaggttataaaaagtgacaactatagtaagcgt |  |
| ara-F1 | tttttataacctccttagagctcgaattcccaaaaaaacg |  |
| ara-R1 | aagcgactgctgctgcaaaattatgacaacttgacggcta |  |
| bb-rha-F | ttttgcagcagcagtcgctt |  |
| tet-bb-R | gaccgctttcgctggagcgcgacga |  |
| tet-F | gcgctccagcgaaagcggtcgtgccacctgacgtctaagaaacca |  |
| tet-pheS-F | gaattcgtcgacaaagaggagaaagatatcatgtcacatctcgcagaactggttg |  |
| tet-R | gatatctttctcctctttgtcgacgaattc |  |
| tet-pheS-R | tctgggtcattttcggcgagcccattcgccaatccggatatagtt |  |
| tet-bb-F | ctcgccgaaaatgacccagagcgctgccg |  |
| fhuA-F | aatataccatgggcagcagcatggcgcgttccaaaactgc | Construction of pET28a-fhuA-mCherry and pET28a-fhuA Δ236-248-mCherry |
| fhuA-R | accatgccgccgccgccgccgccgaaacggaaggttgcgg |  |
| fhuA-248-F | gtcttgcgcgttctgccaatgcaccggcgttcacctggcg |  |
| fhuA-235-R | attggcagaacgcgcaagac |  |
| pet-lac-bb-F | gatccggctgctaacaaagcccgaaaggaagctgagttgg | Cloning of the pET28a backbone and the pET28a-mCherry backbone |
| backbone-F | ggcggcggcggcggcggcatggtgagcaag |  |
| pet-lac-bb-R | gctgctgcccatggtatatt |  |
| lac-YFP-F | aatataccatgggcagcagcatgcgtaaaggagaagaact | Construction of pET28a-pheS-His_6_, pET28a-EcpheS-YFP-His_6,_ and pET28a-YFP-His_6_ |
| YFP-His-R | gctttgttagcagccggatctcagtggtggtggtggtggtgtttgtatagttcatccatg |  |
| pheS-His-R | gctttgttagcagccggatctcagtggtggtggtggtggtgtttaaactgtttgaggaaac |  |
| pheT-His-R | gctttgttagcagccggatctcagtggtggtggtggtggtgatccctcaatgatgcctgga | Construction of pET28a-pheT-His_6_ and pET28a-pheT^D1^-His_6_ |
| pheTD1-His-R | gctttgttagcagccggatctcagtggtggtggtggtggtgggctgcctgtaccggctcat |  |
| mCh-His-R | gctttgttagcagccggatctcaGTGGTGGTGGTGGTGGTGcttgtacagctcgtccatg | Construction of pET28a-mCherry-His_6_, pET28a-His_6_-mCherry-EcdnaA, and pET28a-dnaA-mCherry-His_6_ |
| His-mdA-F | aatataccatgggcagcagcatgCACCACCACCACCACCACgtgagcaagggcgaggagga |  |
| hisorange-F | aatataccatgggcagcagcatgCACCACCACCACCACCACgtgagcaagggcgaggaga | Construction of pET28a-His_6_-mOrange-hslU |
| orangehU-R | tttcgcgtggggtcatttcagagccgccgccgccgccgcccttgtacagctcgtccatgc |  |
| hU-F | tctgaaatgaccccacgcgaaa |  |
| His-hslU-R | gctttgttagcagccggatctcattataggataaaacggctcagat |  |
| 28Kl_057_KPpS_F | aatataccatgggcagcagcATGTCACATCTCGCAGAGCTG | Construction of pET28a-KppheS-YFP-His_6_ and pET28a-BspheS-YFP-His_6_ |
| 28Kl_058_KPpS_CYFP_R | atgccgccgccgccgccgccTTTAAACTGTTTGAGGAAACGCAGATCG |  |
| 28Kl_054_CYFP_F | ggcggcggcggcggcggcatgcg |  |
| 28Kl_053_CmChH_R | gctttgttagcagccggatctcaG |  |
| 28Kl_055_BSpS_F | aatataccatgggcagcagcatggaagaaaagctaaaacagctgg |  |
| 28Kl_056_BSpS_CYFP_R | atgccgccgccgccgccgcccgcctgtttaaactgcgaaataaatctg |  |
| 28Kl_052_NHmCh_F | aatataccatgggcagcagcatgC | Construction of pET28a-His_6_-mCherry-KpdnaA and pET28a-His_6_-mCherry-BsdnaA |
| 28Kl_043_Nmch_R | gccgccgccgccgccgcccttgtacag |  |
| 28Kl_050_KPdA_Nmch_F | agggcggcggcggcggcggcTCACTTTCGCTTTGGCAGCAG |  |
| 28Kl_051_KPdA_R | gctttgttagcagccggatcTCACGAGGATAATGTTCTGATTAAATTGG |  |
| 28Kl_046_BSdA_Nmch_F | agggcggcggcggcggcggcGAAAATATATTAGACCTGTGGAACC |  |
| 28Kl_047_BSdA_R | gctttgttagcagccggatcCTATTTAAGCTGTTCTTTAATTTCTTTTACATGC |  |
| DI_pheT_N_fw | agcctgatgggataggctctaagtccaacgaaccagtgtcgtaggctggagctgcttcg | λ Red of *pheT* upstream for phage transduction |
| DI_pheT_N_rv | acggcgataaaagtcaatgttttgatggcgttgaaacgaagatgccgggagcagacaag |  |
| Ci_hslVU_fw | tgcgttggtggcagatgaagatctgagccgttttatcctatgtaggctggagctgcttc | λ Red of *hslVU-ECFP* |
| Ci_hslVU_rv | agccccatcaaacaatgatgaaaatgattgaacgcgattagatgccgggagcagacaag |  |
| CiECFP_hslVU_fw | tggtggcagatgaagatctgagccgttttatcctaggcggcggcggcggcggcatgctg |  |
| CiECFP_hslVU_rv | ccccatcaaacaatgatgaaaatgattgaacgcgattacttgtacagctcgtccatgcc |  |
| Ci_dnaA_fw | agaagatttttcaaatttaatcagaacattgtcatcgtaatgtaggctggagctgcttc | λ Red of *dnaA-mCherry, mCherry-dnaA*, and *dnaA-mCherry-In* |
| Ci_dnaA_rv | ttttaataaatgctcacgttctacggtaaatttcataggtgatgccgggagcagacaag |  |
| Ni_dnaA_fw | ttagatcgattaagccaatttttgtctatggtcattaaatgtaggctggagctgcttcg |  |
| Ni_dnaA_rv | ctgccgcgaggcgggcacgatttacgccgcatattggaaagatgccgggagcagacaag |  |
| dnaA_dn | ggcagaccacggcagatatc |  |
| dnaA_Nimch_fw | gtacgacctcacaccagtgg |  |
| dnaA_Nimch_rv | caaaacggtttggcgcgtac |  |
